# SARS-CoV-2 spike-induced syncytia are senescent and contribute to exacerbated heart failure

**DOI:** 10.1101/2022.10.10.511541

**Authors:** Luming Wan, Huilong Li, Muyi Liu, Enhao Ma, Linfei Huang, Yilong Yang, Qihong Li, Yi Fang, Jingfei Li, Bingqing Han, Chang Zhang, Lijuan Sun, Xufeng Hou, Haiyang Li, Mingyu Sun, Sichong Qian, Xuejing Duan, Ruzhou Zhao, Xiaopan Yang, Yi Chen, Shipo Wu, Xuhui Zhang, Yanhong Zhang, Gong Cheng, Gengye Chen, Qi Gao, Junjie Xu, Lihua Hou, Congwen Wei, Hui Zhong

## Abstract

Patients with pre-existing heart failure are at a particularly high risk of morbidity and mortality resulting from SARS-CoV-2 infection. Direct acute cardiac injury or cytokine storms have been proposed to contribute to depressed cardiac function. However, the pathogenic mechanisms underlying the increased vulnerability to heart failure in SARS-CoV-2 infected patients are still largely unknown. Here, we found that the senescent outcome of SARS-CoV-2 spike protein (SARS-2-S)-induced syncytia exacerbated heart failure progression. We first demonstrated that syncytium formation in cells expressing SARS-2-S delivered by DNA plasmid or LNP-mRNA exhibits a senescence-like phenotype. Extracellular vesicles containing SARS-2-S (S-EVs) also confer a potent ability to form senescent syncytia without *denovo*synthesis of SARS-2-S. Mechanistically, SARS-2-S syncytia provoke the formation of functional MAVS aggregates, which regulate the senescence fate of SARS-2-S syncytia by TNF α . We further demonstrate that senescent SARS-2-S syncytia exhibit shrinked morphology, leading to the activation of WNK1 and impaired cardiac metabolism. In pre-existing heart failure mice, the WNK1 inhibitor WNK463, anti-syncytial drug niclosamide, and senolytic dasatinib protect the heart from exacerbated heart failure triggered by pseudovirus expressing SARS-2-S (SARS-2-Spp). Signs of senescent multinucleated cells are identified in ascending aorta from SARS-CoV-2 omicron variant-infected patient. Our findings thus suggest a potential mechanism for COVID-19-mediated cardiac pathology and recommend the application of WNK1 inhibitor for therapy.

**Significance Statement:** In this paper, we directly linked SARS-2-S-triggered syncytium formation with the ensuing induction of cellular senescence and its pathophysiological contribution to heart failure. We propose that both SARS-2-S expression and SARS-2-S protein internalization were sufficient to induce senescence in nonsenescent ACE2-expressing cells. This is important because of the persistent existence of SARS-2-S or extracellular vesicles containing SARS-2-S during the acute and post-acute stages of SARS-CoV-2 infection in human subjects. In searching for the underlying molecular mechanisms determining syncytial fate, the formation of functional MAVS aggregates dependent on RIG-I was observed at an early stage during fusion and regulated the anti-death to senescence fate of SARS-2-S syncytia through the TNFα-TNFR2 axis. We also found impaired cardiac metabolism in SARS-2-S syncytia induced by condensed WNK1. Importantly, SARS-2-Spp-exacerbated heart failure could be largely rescued by WNK1 inhibitor, anti-syncytial drug or senolytic agent. Together, we suggest that rescuing metabolism dysfunction in senescent SARS-2-S syncytia should be taken into consideration in individuals during the acute or post-acute stage of SARS-CoV-2 infection.

## Introduction

Coronavirus disease 2019 (COVID-19), caused by severe acute respiratory syndrome coronavirus 2 (SARS-CoV-2), has resulted in a global pandemic with millions of deaths^1^. Although symptoms resulting from infection typically resolve within weeks, some individuals experience persistent symptoms following the acute phase of COVID-19, the so-called post-acute sequelae of COVID-19 (PASC) or long COVID^2–4^. SARS-CoV-2 infection predominantly offends the respiratory system. Currently, evidence has suggested cardiac complications as one of the important pathogenic features of COVID-19^5, 6^. More importantly, compared with patients without heart failure, those with diagnosed heart failure experienced a longer length of hospital stay, increased risk of mechanical ventilation and intensive care unit admission, and higher mortality^7, 8^. Direct myocardial injury, inflammation and thrombosis, hypoxemia, and ACE2 downregulation are suggested to be involved in COVID-19-mediated cardiac pathology; however, the pathogenic mechanisms underlying the increased vulnerability to heart failure in patients with COVID-19 are still largely unknown^9–11^.

SARS-CoV-2 syncytium constitutes a hallmark of COVID-19-associated pathology and may potentially contribute to pathology by facilitating viral dissemination, cytopathy, immune evasion, and the inflammatory response^12^. Syncytium formation is mediated by the SARS-CoV-2 spike protein (SARS-2-S). SARS-2-S interacts with the ACE2 receptor on the host cell and is then proteolytically activated by host proteases, leading to the fusion of the viral membrane with the plasma membrane^13^. In addition to mediate viral-cell fusion, the presence of SARS-2-S on the cell surface is sufficient to mediate cell-cell fusion, a mechanism called fusion from within (FFWI)^14, 15^. The fusogenic activity of SARS-2-S is potent, as even undetectable amounts of SARS-2-S can cause cell‒ cell fusion^14^. In addition to mediating FFWI, virus-like particles or lipid vesicles containing surface SARS-2-S can serve as a bridge between two cells and induce fusion from cells without *de novo* synthesis of SARS-2-S protein (FFWO)^14, 15^. Cell-cell fusion in uninfected cells may also occur in long COVID-19 syndrome, as SARS-2-S or circulating extracellular vesicles containing SARS-2-S (S-EVs) were detected predominantly in PASC patients for up to 12 months after diagnosis^16, 17^.

SARS-CoV-2 elicits the senescence of infected cells, similar to other viruses (virus-induced senescence, VIS)^18, 19^. Cellular senescence is a state of stable cell cycle arrest characterized by increased lysosomal activity, exemplified by induction of the lysosomal enzyme senescence-associated β-galactosidase (SA-β-gal) and activation of the senescence-associated secretory phenotype (SASP)^20–22^. SARS-CoV-2 infection triggers paracrine senescence and can lead to a hyperinflammatory environment and the onset of acute respiratory disease syndrome^23^. It is not entirely clear whether virus propagation stress or specific viral molecular components induce VIS. Interestingly, isolated recombinant SARS-2-S protein has been shown to increase the SASP in senescent ACE2-expressing cells^18^. However, the direct linkage of SARS-2-S syncytia with senescence and the degree to which SARS-2-S syncytia affect pathology in the setting of cardiac dysfunction are unknown.

Here, we demonstrated that both SARS-2-S expression and SARS-2-S protein internalization were sufficient to induce senescence in nonsenescent ACE2-expressing cells. The formation of functional MAVS aggregates dependent on RIG-I was observed at an early stage during fusion and regulated the anti-death to senescence fate of SARS-2-S syncytia through the TNFα-TNFR2 axis. Except for SASP, shrinking of SARS-2-S syncytia triggered WNK1 activation, which led to impaired cardiac metabolism. SARS-2-Spp exacerbated heart failure progression, a phenotype that could be largely rescued by WNK1 inhibitors, anti-syncytial drugs or senolytic agents. Together, we suggest that rescue metabolism dysfunction of senescent SARS-2-S syncytia should be taken into consideration in individuals with a history of heart failure during the acute or post-acute stage of SARS-CoV-2 infection.

## Results

### SARS-2-S and its variant-expressing syncytia exhibit a senescence-like phenotype

Recombinant SARS-2-S protein induces senescence in ACE2-expressing cells. To explore whether SARS-2-S expression was sufficient to induce senescence in nonsenescent ACE2-expressing cells, we cocultured SARS-2-S (Wuhan-Hu-1)-expressing A549 cells with ACE2-expressing A549 cells at a 1:1 ratio (Extended Data Fig. 1a-c). Syncytial cells rapidly appeared at 4 h post coculture (hpc) and gradually grew in size, as indicated by microscopic analysis (Fig. 1a). Notably, SARS-2-S-expressing syncytia (hereafter referred to as SARS-2-S syncytia) with enhanced SA-β-gal staining began to appear at 12 hpc, and the percentage of SA-β-gal-positive syncytia among SARS-2-S syncytia gradually increased during the fusion process (Fig. 1a). In contrast to SARS-2-S, a SARS-2-S_NF_ mutant (R682A R683A R685A K814A R815A), which fails to cause fusion because it cannot be cleaved by host protease, lost the ability to induce senescence^24, 25^ (Fig. 1a, Extended Data Fig. 1d). We did not observe multinucleated cells or SA-β-gal–positive staining in A549 cells that only expressed SARS-2-S or ACE2 at the time points tested (Extended Data Fig. 1e). Quantitative real-time PCR (RT-qPCR) revealed significant transcriptional upregulation of senescence-associated genes (Fig. 1b). Consistently, the conditioned medium of SARS-2-S syncytia at 48 hpc contained significantly more SASP-related cytokines, such as IL-6, IL-8, and IL-2, than that at 0 hpc (Fig. 1c). In a similar manner, we observed SARS-2-S–induced senescence during coculture of the same and different cell types (Fig. 1d). In A549 cocultured cells, the appearance of SA-β-gal-positive staining was associated with the syncytial nuclei number. Eventually, over 70% of SARS-2-S syncytia exhibited positive SA-β-gal staining at 72 hpc (Fig. 1e). We next compared the senescence potential of four Omicron variants, SARS-2-S_BA.1_, SARS-2-S_BA.5_, SARS-2-S_BA.2.75_, and SARS-2-S_BA.2.75.2_, with that of SARS-2-S (Extended Data Fig. 1d). As reported, the Omicron BA.1 variant showed significantly weaker fusogenic activity than SARS-2-S^26^ (5× reduction). The fusogenic capacity of BA.5 (1.2× reduction), BA.2.75 (1.7× reduction), and BA.2.75.2 (1.7× reduction) was modestly lower than that of SARS-2-S^27, 28^ (Extended Data Fig. 1f). Notably, the percentage of SA-β-gal-positive syncytia was comparable to that of SARS-2-S, although they formed more slowly (Fig. 1f, g). Furthermore, SARS-2-S syncytia could be selectively killed by senolytic drugs, including the multiple-kinase inhibitor fisetin, the tyrosine kinase inhibitor dasatinib, and the BCL-2–specific venetoclax (Fig. S1g). Taken together, the above observations indicate that SARS-2-S elicits a senescence-like phenotype in an FFWI manner.

**Fig. 1.**
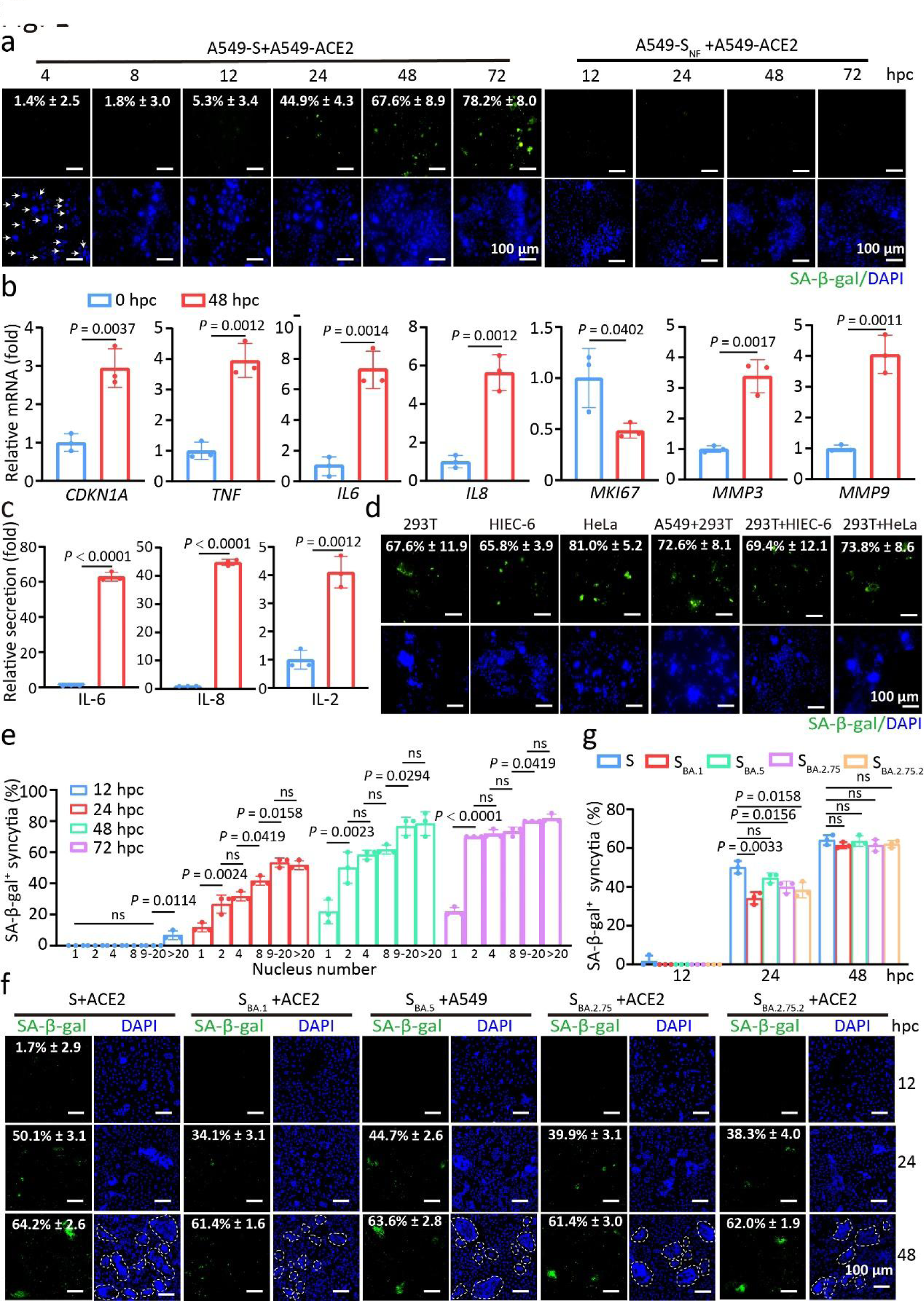
SARS-2-S–expressing syncytia exhibit a senescence-like phenotype. a, SA-β -gal staining of A549-SARS-2-S (SARS-2-S hereafter referred to S) or A549-S_NF_ (R682A, R683A, R685A, K814A, and R815A mutant) cocultured with A549-ACE2 cells for the indicated h (hpc). Green, SA-β-gal staining; blue, nuclear DAPI staining. Scale bars represent 100 μ m. b, Normalized expression of *CDKN1A*, *TNF*,*IL6*,*IL8*, *MKI67*, *MMP3*and *MMP9*transcripts in cocultured A549 cells at 48 hpc relative to that at 0 hpc by RT-qPCR. c, IL-6, IL-8, and IL-2 protein levels in conditioned medium from cells in b by ELISA. d, SA-β-gal staining of S–expressing cells cocultured with the same or different cell types. Scale bars represent 100 μ m. e, Quantification of SA-β-gal positivity in cocultured A549 cells with different nuclei numbers during fusion for the indicated hpc. f, g, SA-β-gal staining (f) SA-β-gal-positivity quantification (g) of S, Omicron variant BA.1 spike (S_BA.1_), Omicron variant BA.5 spike (S_BA.5_), Omicron variant BA.2.75 spike (S_BA.2.75_), and Omicron variant BA.2.75.2 spike (S_BA.2.75.2_)-transfected A549 and A549-ACE2 cells cocultured for the indicated hpc. Scale bars represent 100 μm. All images are representative of 3 biological replicates. All quantified data are presented as the mean ± SD of n = 3 independent experiments. Statistical significance was determined using two-tailed Student’s t test (b, c) or one-way ANOVA and Bonferroni’s post hoc analysis (e, g).

### SARS-2-S delivery by EVs or mRNA confers the ability to trigger syncytial senescence

Since SARS-2-S was shown to be incorporated into extracellular vesicles circulating in patient blood^29^, we then asked if SARS-2-S protein internalization by S-EVs has the ability to trigger syncytium formation and cell senescence in an FFWO-dependent manner. To this end, we first used ultracentrifugation to purify EVs from the supernatant of HeLa-or SARS-2-S-expressing HeLa cells and confirmed their identity by probing for exosome markers (Extended Data Fig. 2a, b). We next labelled these EVs with the fluorescent dye DiO and exposed A549 cells to the EVs, and the expression of SARS-2-S on the surface of the cells that took up the S-EVs was observed (Extended Data Fig. 2c, d). These cells fused with cells expressing ACE2 at 8 hpc, as indicated by the appearance of multinucleated cells (Fig. 2a). Importantly, the fused cells exhibited enhanced SA-β-gal and p21 staining, with increased transcription of the *TNF*,*IL6*,*IL8*, and *CDKN1A*genes (Fig. 2b), features that are typical of senescence. We also noted the senescence phenotype in other similarly treated cell types, but not in cells that took up EVs lacking SARS-2-S (Extended Data Fig. 2e, f). In addition, A549-ACE2 cells were incubated with S-EVs directly for 2 to 48 h. Notably, S-EVs induced syncytium formation as early as 2 h, a timeframe in which adjacent cells may fuse directly without S-EVs uptake and SARS-2-S presentation at the cell surface. The appearance of senescent syncytia began to occur at 12 h, (Fig. 2c, d). Eventually, approximately 70% of syncytia exhibited positive SA-β-gal staining at 48 h upon S-EVs addition (Fig. 2c). Therefore, SARS-2-S protein internalization was sufficient to induce senescence in an FFWO manner.

**Fig. 2.**
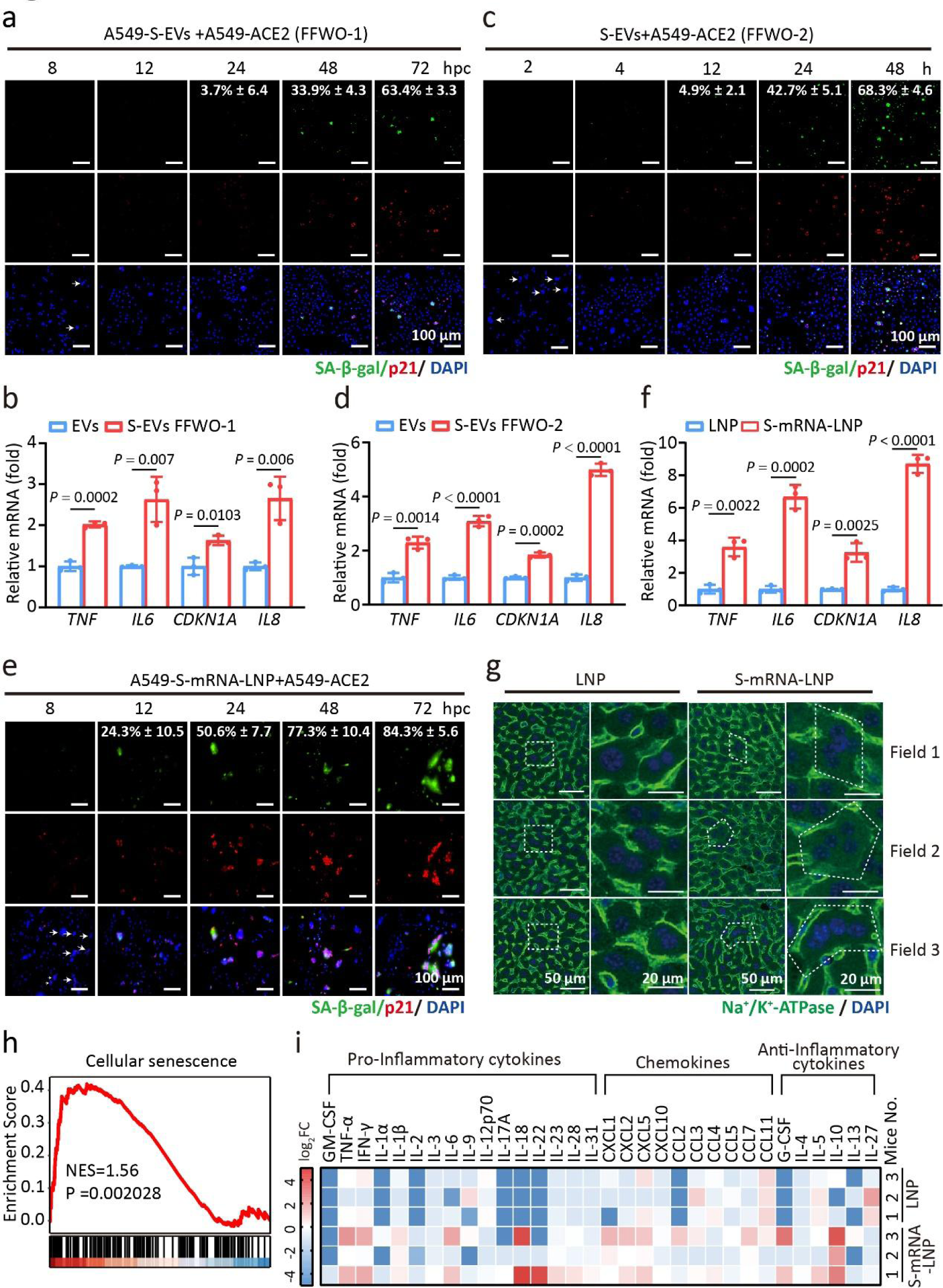
SARS-2-S delivery by EVs or mRNA triggers syncytial senescence. a, c, e, SA-β-gal and p21 staining of A549-S-EVs and A549-ACE2 cocultured cells (a, fusion from without (FFWO-1)) or A549-ACE2 cells treated with S-EVs (c, FFWO-2) or A549-S-mRNA-LNP and A549-ACE2 cocultured cells (e) for the indicated times. Green, SA-β-gal staining; red, p21 staining; blue, DAPI staining. Scale bars represent 100 μm. b, d, f, Normalized expression of *TNF*,*IL6,CDKN1A, andIL8*in A549 cells at 24 hpc from a (b), c (d), and e (f) relative to that at 0 hpc by RT-qPCR. g, Immunohistochemical analysis of liver tissues from mice intramuscularly injected with a single dose of S-mRNA-LNP (0.12 mg/kg) or LNP at 24 h after injection. Green, Na^+^/Ka^+^-ATPase staining of the cell membrane; blue, DAPI staining. Scale bars represent 50 μm or 20 μm as indicated. h, Gene set enrichment analysis (GSEA) of genes related to cellular senescence in the livers from mice in g. NES, normalized enrichment score. i, Multiplex bead-based protein analysis in the serum from mice in g. All images are representative of 3 biological replicates. All quantified data are presented as the mean ± SD of n = 3 independent experiments. Statistical significance was determined using two-tailed Student’s *t*test (b, d, f).

We next asked whether SARS-2-S delivered in the form of encoding mRNA would also trigger syncytium formation and cell senescence (Extended Data Fig. 3a). To this end, we packaged the encoding mRNA into a lipid nanoparticle formulation (S-mRNA-LNP). Western blotting confirmed the expression of SARS-2-S treated with S-mRNA-LNP (Extended Data Fig. 3b). Next, we cocultured S-mRNA-LNP-treated A549 cells with A549-ACE2 cells at a 1:1 ratio. Notably, syncytial cells formed rapidly and began to stain positive for SA-β-gal and p21 at 8 hpc (Fig. 2e). Significant upregulation of senescence-associated genes supported the presence of a senescence phenotype (Fig. 2f). We obtained similar results using 293T-ACE2 cells, but did not observe obvious signs of senescence in LNP- or S-mRNA-LNP-treated A549 cells (Extended Data Fig. 3c-e). This finding indicates that SARS-2-S has the ability to trigger syncytial senescence regardless of its manner of delivery.

Finally, we explored whether cell senescence related to SARS-2-S-mediated cell fusion would also occur*invivo*. To this end, we injected AAV-hACE2 constructs into the tail veins of 8-week-old C57BL/6J mice to drive mainly hepatic hACE2 expression (Extended Data Fig. 3f, g). Four weeks later, we intramuscularly administered a single dose of S-mRNA-LNP or LNP to hACE2-expressing mice and examined the induction of senescence. Immunohistochemical analysis revealed robust SARS-2-S expression and the increased appearance of multinucleated cells with at least four nuclei in the livers of S-mRNA-LNP-treated mice compared to the livers of LNP-treated mice (Extended Data Fig. 3h and Fig. 2g). RNA sequencing (RNA-seq) of liver samples revealed the upregulated expression of cellular senescence-related genes (Fig. 2h). KEGG pathway analysis of differentially expressed genes (DEGs) revealed enrichment cellular senescence pathways (Extended Data Fig. 4). Finally, multiplex protein analysis of serum samples from these animals revealed that S-mRNA-LNP strongly induced SASP-reminiscent cytokines (Fig. 2i). We therefore concluded that SARS-2-S delivery in the form of DNA, EVs, or mRNA can trigger the formation of syncytia with a senescence phenotype.

### SARS-2-S syncytia provoke the formation of functional MAVS aggregates

Cell-cell fusion in the absence of SARS-CoV-2 infection activates the cGAS-STING pathway, which has been suggested to promote cellular senescence^30^. However, the occurrence of senescence in SARS-2-S syncytia formed by 293T cells lacking STING expression^31^ indicated the existence of other signalling pathways that regulate the senescence fate of SARS-2-S syncytia. We then focused on mitochondrial antiviral signalling adaptor protein (MAVS), a mitochondrial adaptor protein that links the cytoplasmic RNA sensor RIG-I to its downstream signalling molecules by forming well-ordered prion-like aggregates^32^. As reported, the MAVS staining pattern became noticeably speckled in response to vesicular stomatitis virus (VSV) infection^33^ (Fig. 3a). Strikingly, MAVS was also redistributed as early as 3 hpc in SARS-2-S syncytia in a manner similar to its distribution in mitochondria (Fig. 3a and Extended Data Fig. 5a). Furthermore, MAVS aggregation during cell fusion proceeded in a manner similar to that induced by VSV infection, as determined by semidenaturing detergent agarose gel electrophoresis (SDD-AGE) (Fig. 3b). While VSV infection began to trigger massive mitochondrial elongation and tubularization at 6 h post infection (hpi), we observed enhanced speckled staining of the mitochondria in SARS-2-S syncytia, with relatively less mitochondrial fusion, during the entire fusion process (Fig. 3a). Importantly, MAVS aggregation was reduced by knockdown of mitofusin 1 (MFN1) but not by knockdown of OPA1; both of these proteins are mitochondrial membrane proteins that regulate mitochondrial dynamics and RIG-I-induced antiviral signalling^34^ (Fig. 3c and Extended Data Fig. 5b). Accordingly, we knocked down the RIG-I gene (Extended Data Fig. 5b). We found that this significantly reduced MAVS aggregation in cocultured cells (Fig. 3d). MAVS aggregation in cocultured cells was also reduced by MDA5 silencing, but this effect was much less pronounced than that observed upon RIG-I silencing, although the knockdown efficiency was approximately 70% for both genes (Fig. 3d and Extended Data Fig. 5b). These data suggest the involvement of RIG-I, the primary sensor of nonself-derived dsRNA, in functional MAVS aggregation in SARS-2-S syncytia.

**Fig. 3.**
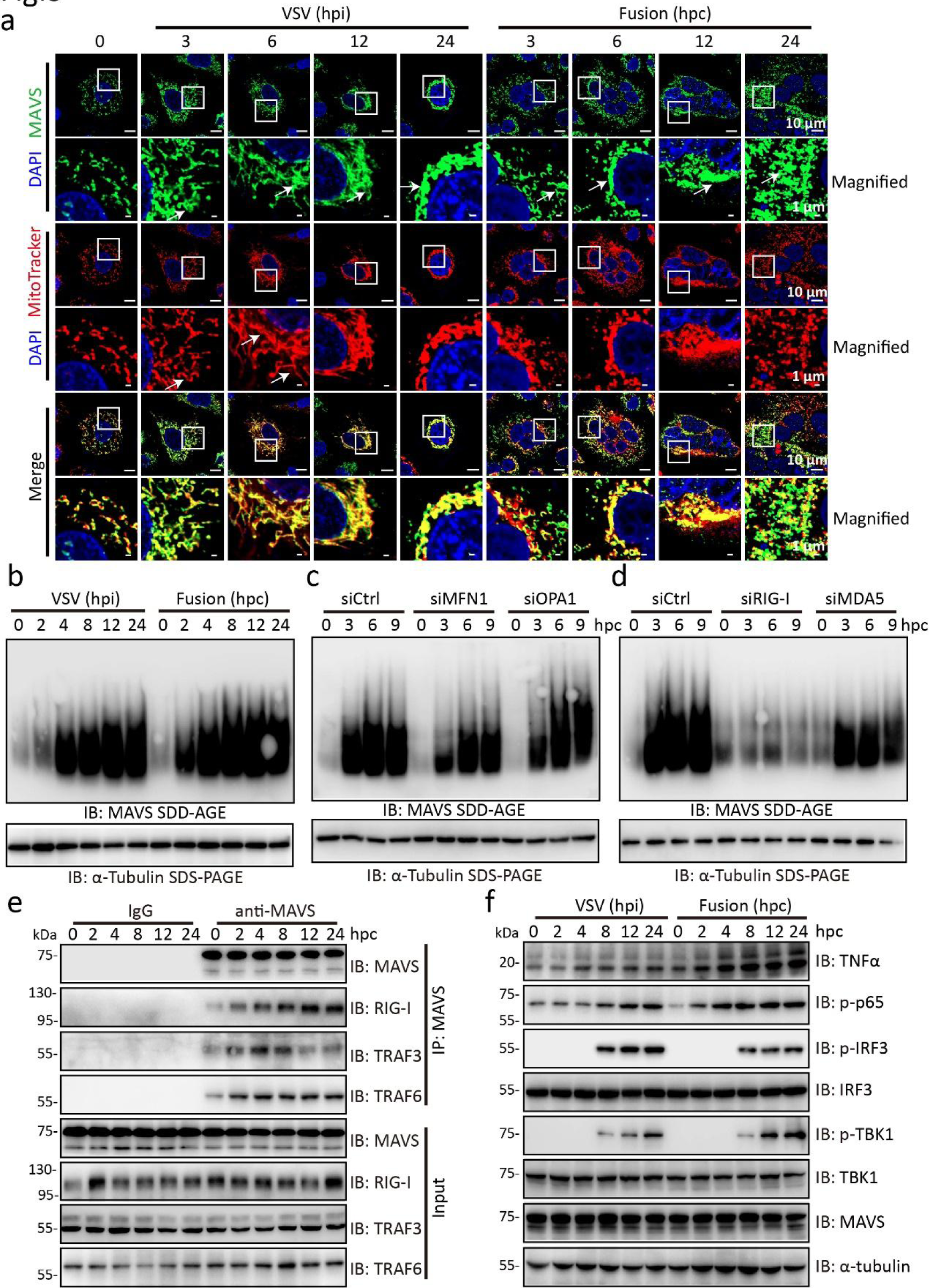
SARS-2-S syncytia induce the RIG-I–dependent formation of functional MAVS aggregates. a, Confocal microscopic images of MAVS and mitochondria in A549 cells infected with VSV or A549 cells cocultured for the indicated times. Red, MitoTracker staining for mitochondria; green, MAVS staining; blue, DAPI staining. A magnified view of the boxed region is also shown. Scale bars represent 10 μ m (whole image) and 1 μ m (magnified). b, MAVS aggregation in VSV-infected A549 cells or A549 cells cocultured for the indicated times by SDD-AGE with an anti-MAVS antibody. SDS-PAGE immunoblotting of α -tubulin was used as a loading control. c, d, MAVS aggregation in sicontrol (siCtrl), siMFN1, and siOPA1 (c) or siCtrl, siRIG-I, and siMDA5 (d) A549 cells cocultured for the indicated times. All these siRNA oligos were transfected into A549-S and A549-ACE2 cells 24 h before the start of coculture. α-Tubulin was used as the loading control. MAVS aggregates were analysed by SDD-AGE as in b. e, Immunoprecipitation analysis of the MAVS interaction with RIG-I, TRAF3, and TRAF6 in A549 cells cocultured for the indicated times. f, Immunoblotting analysis of proteins of the activated IFN-I signalling pathway in VSV-infected A549 cells or A549 cocultured cells for the indicated times.α-Tubulin was used as the loading control. All images are representative of three independent experiments.

To confirm the spatiotemporal events linking RIG-I and MAVS during fusion, we performed a kinetic analysis of MAVS-RIG-I complex formation during cell fusion. Following the immunoprecipitation of endogenous MAVS, we detected increased levels of the RIG-I, TRAF3, and TRAF6 proteins in the immunoprecipitates as early as 2 hpc (Fig. 3e). In particular, the fusion process was accompanied by increased phosphorylation levels of p65, IRF3, and TBK1, a measure of IFN-I signalling activity (Fig. 3f). Finally, the activity of the NF-κB promoter was dramatically enhanced in cocultured cells (Extended Data Fig. 5c). The above observations indicate the activation of RIG-I-MAVS signalling in SARS-2-S syncytia.

### MAVS-TNFα signalling regulates the senescence fate of SARS-2-S syncytia

We next attempted to identify factors that determine the senescence fate of SARS-2-S syncytia. To this end, we systematically analysed the gene expression of cocultured cells by RNA-seq at 4 and 48 hpc. Principal component analysis (PCA) of the data revealed that samples obtained after coculture for these durations did not overlap and were well separated from their respective controls (Fig. 4a). Gene Ontology (GO) enrichment analysis indicated that the coverage of senescence- and SASP-related GO terms increased with increasing coculture time (Fig. 4b). This finding supported the notion that fusogenic SARS-2-S gradually triggers a senescence stress response. Importantly, KEGG pathway analysis revealed enrichment in the TNFα and NF-κB signalling pathways throughout the fusion process (Fig. 4c and Extended Data Fig. 6a, b). Notably, MAVS or RIG-I silencing significantly reduced the activation of NF-κB, IRF3, and TNFα (Fig. 4d). Furthermore, the expression of TNFα and IL6 was also largely abolished in MAVS KO-, siRIG-I-, or siMAVS-cocultured 293T cells (Fig. 4e, f). In A549 cells with intact cGAS-STING signalling, syncytial survival and the senescent syncytial subpopulation were also significantly reduced by MAVS or RIG-I knockdown, but not by STING knockdown (Fig. 4g, h). These results indicated the essential role of RIG-I-MAVS in regulating the senescence fate of the SARS-2-S syncytium.

**Fig. 4.**
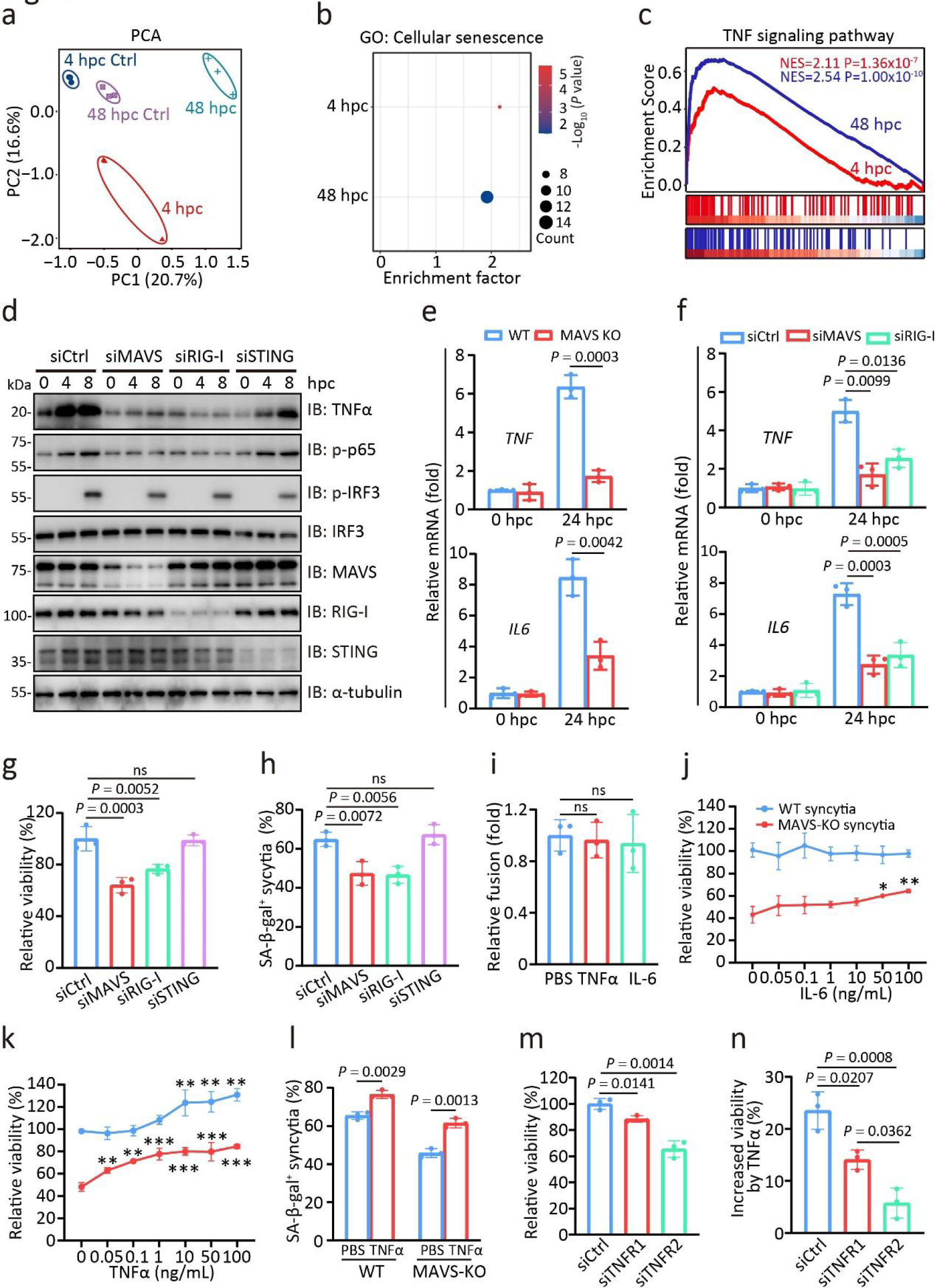
MAVS–TNFα signalling regulates the survival to senescence fate of SARS-2-S syncytia. a, PCA of gene expression patterns in A549 cells cocultured for the indicated times. In all plots, each point represented an independent biological replicate. b, Senescence pathways enriched in the samples analysed in a based-on GO term analysis. c, GSEA of genes related to the TNF signalling pathway among the DEGs in cells from a. d, Immunoblotting analysis of proteins of the activated IFN-I signalling pathway in siCtrl, siMAVS, siRIG-I, or siSTING cells cocultured for the indicated times. All these siRNA oligos were transfected into A549-S and A549-ACE2 cells 24 h before the start of coculture. α-Tubulin was used as the loading control. e, f, Normalized expression of *TNF*and*IL6*in WT and MAVS-KO cocultured 293T cells (e) or siCtrl, siMAVS, and siRIG-I (f) cocultured 293T cells at 24 hpc relative to that at 0 hpc. g, Relative cell viability of siMAVS, siRIG-I, or siSTING cocultured cells at 24 hpc by CCK8 assay. The cell viability of siCtrl cocultured cells at 24 hpc was set to 100%. All these siRNA oligos were transfected into A549-S and A549-ACE2 cells 24 h before coculture. h, SA-β-gal positivity in cocultured cells from g. i, Relative fusion of cocultured A549 cells at 24 hpc in the presence of TNFα (100 ng/mL) or IL-6 (100 ng/mL) by β-galactosidase assay. The fusion level of cocultured A549 cells treated with PBS was set to 1. j, k, Relative cell viability of WT or MAVS-KO cocultured cells at 48 hpc in the presence of IL-6 (j) or TNFα (k). The cell viability of WT cocultured cells at 48 hpc was set to 100%. l, SA-β-gal positivity in WT or MAVS-KO cocultured cells at 48 hpc in the presence of TNF α . m, Relative viability of siTNFR1- or siTNFR2-cocultured A549 cells at 24 hpc. The viability of siCtrl cells at 24 hpc was set to 100%. n, Increased cell viability of siCtrl, siTNFR1, or siTNFR2 cocultured cells at 48 hpc in the presence of TNFα than that in the presence of PBS. All quantified data are shown as the mean ± SD of n = 3 independent experiments. Statistical significance was determined using two-tailed Student’s *t* test (e) or one-way ANOVA and Bonferroni’s post hoc analysis (f, g, h, i, j, k, l, m, n).

To determine whether MAVS-mediated NF-κB signalling induction is required for the survival and senescence of SARS-2-S syncytia, we supplemented WT and MAVS KO cocultured 293T cells with increasing concentrations of recombinant human TNFα and IL-6. While pretreatment with TNFα and IL-6 before coculture did not affect SARS-2-S fusogenic potential (Fig. 4i), recombinant IL-6 at concentrations over 50 ng/mL improved the viability of MAVS KO cocultured cells, and recombinant TNFα at concentrations above 0.05 ng/mL improved the viability of MAVS KO cocultured cells (Fig. 4j, k and Extended Data Fig. 7a, b). Notably, the percentage of SA-β-gal-positive syncytia was also significantly increased by TNFα, especially in MAVS KO cells (Fig. 4l). TNFα exerts its activity by stimulating its receptors, TNFR1 and TNFR2, which trigger distinct and common signalling pathways that control cell apoptosis and survival^35–37^. We observed that TNFR2 *(TNFRSF1B)*was selectively upregulated in cocultured A549 cells at both the early and late stages of the fusion process (Extended Data Fig. 7c). Indeed, knockdown of TNFR1 and TNFR2 significantly reduced the viability of cocultured cells (Fig. 4m and Extended Data Fig. 7d). Remarkably, TNFR1 knockdown blocked the increase of cocultured cell viability induced by TNFα, but it was not as effective as TNFR2 knockdown (Fig. 4n). Furthermore, neither TNFR1 nor TNFR2 deficiency affected SARS-2-S fusogenicity (Extended Data Fig. 7e). Taken together, these observations suggest that the MAVS-TNFα-TNFR2 axis regulates the anti-death to senescence fate of SARS-2-S syncytia.

### Senescent SARS-2-S syncytia shrinkage triggers WNK1 phase separation

A defining feature of cellular senescence is the enlargement and vacuolization of the cell body^38^. Here in our assay, the size of A549 senescent cells induced by the established senescence stressor palbociclib for 7 days did not differ from control cells treated with PBS (Fig. 5a). However, we noticed distinct morphological changes during SARS-2-S-triggered fusion progression. SARS-2-S syncytia of A549 began to exhibit a smaller surface area per nucleus regardless of the nuclei numbers with a smaller size tendency at 24 hpc (Fig. 5b, c). The disproportionate decrease in cell volume per single nucleus was also consistently evident in senescent cardiomyocyte AC16 syncytia (Fig. 5d). Importantly, these SA-β-gal-positive syncytia were alive for at least 7 observational days after coculture (Extended Data Fig. 8a, b). The shrinkage of senescent SARS-2-S syncytia made us wonder whether with-no-lysine (WNK) kinase1, a molecular crowding sensor that undergoes cell shrinkage-dependent phase separation^39^, was activated. Notably, we observed a significant shift of WNK1 from a diffuse to punctate distribution in senescent SARS-2-S syncytia of A549 and AC16 cells (Fig. 5e-i), but not in those cells treated with palbociclib (Fig. 5j-m). This phase behaviour appeared as early as 12 hpc in AC16 cells (Fig. 5e). Accordingly, WNK1-triggered SPAK phosphorylation began to appear at 24 hpc and persisted for 72 hpc (Fig. 5n-p). No signs of SPAK phosphorylation signals were observed in SARS-2-SNF cocultured cells (Fig. 5o, p). Therefore, senescent SARS-2-S syncytia exhibit shrinked morphology, leading to the activation of WNK1.

**Fig. 5.**
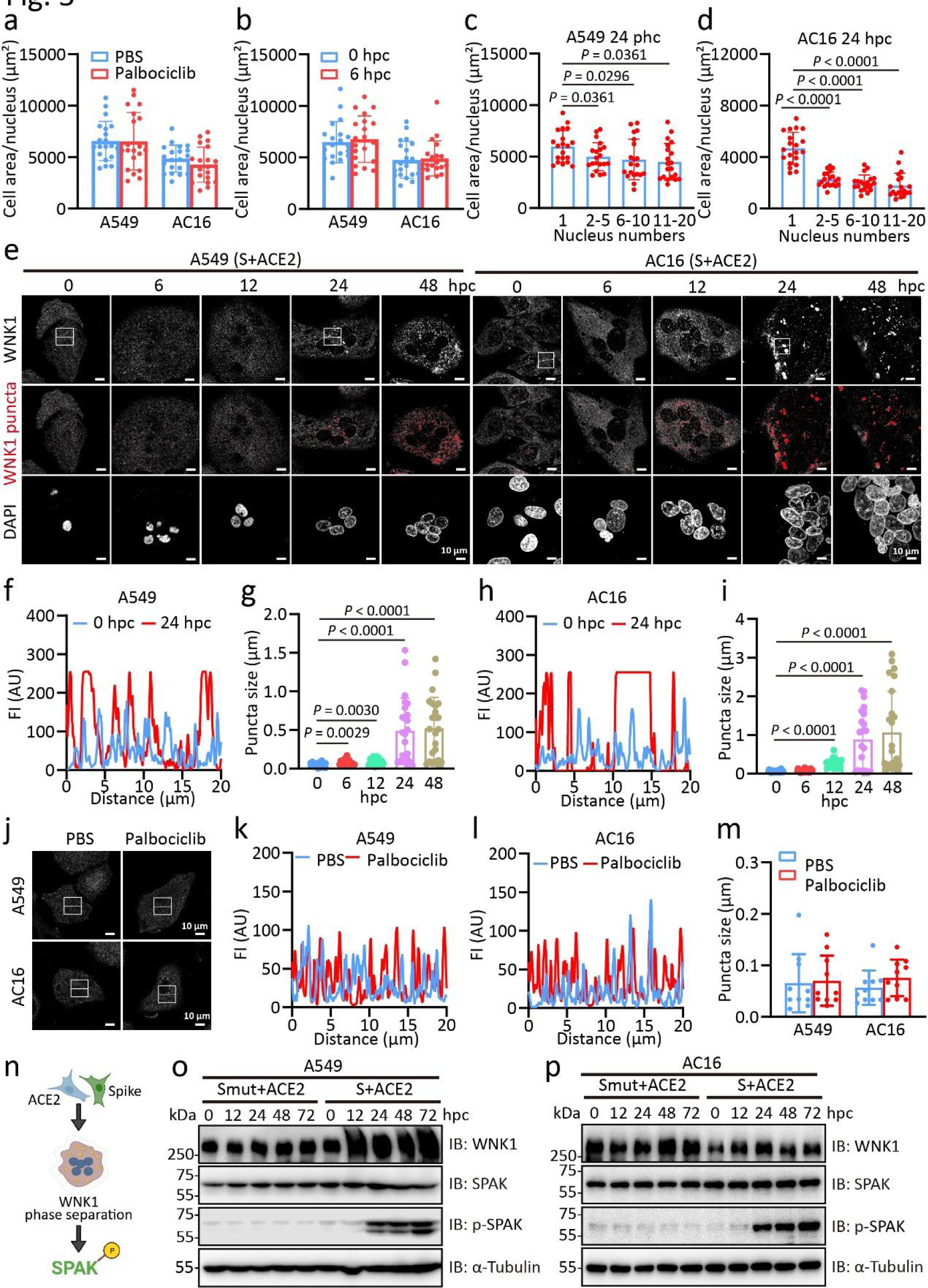
Senescent SARS-2-S syncytia shrinkage triggers WNK1 phase separation. a, Quantification of cell area per nucleus of A549 and AC16 cells treated with palbociclib (1 μM) for 7 days (n = 20 cells). b, Quantification of cell area per nucleus in cocultured A549 or AC16 cells for 6 hpc (n = 20 cells). c-d, Quantification of cell area per nucleus in cocultured A549 cells (c) or AC16 cells (d) with 1, 2-5, 6-10, and 11-20 nuclei numbers at 24 hpc (n = 20 cells). e, Confocal microscopic images of WNK1 in cocultured A549 or AC16 cells for the indicated times. White, WNK1 staining; red, WNK1 puncta. Scale bars represent 10 μm. f-g, Line scans represent fluorescence intensities (FIs) of the inset arrows (f) and quantification of average high-intensity puncta size/confocal field (n = 20 fields) (g) of A549 cells in e. h-i, Line scans represent FIs of the inset arrows (h) and quantification of average high-intensity puncta size/confocal field (n = 20 fields) (i) of AC16 cells in e. The high-intensity threshold signal was kept constant across all conditions analysed. j, Confocal microscopic images of WNK1 in A549 or AC16 cells treated with PBS or palbociclib (1 μM) for 7 days. White, WNK1 staining. k-l, Line scans represent FIs of the inset arrows in j from A549 (k) or AC16 (l) cells. The high-intensity thresholded signal was kept constant across all conditions analysed. m, Quantification of average high-intensity puncta size/confocal field in j (n = 10 fields). n, Scheme of phosphorylation-dependent regulation of the SPAK by WNK1. o-p, Immunoblotting analysis of WNK1, SPAK, and SPAK phosphorylation in S or SNF cocultured A549 (o) or AC16 (p) cells for the indicated times. α-Tubulin was used as the loading control. All images are representative of 3 biological replicates. All quantified data are presented as the mean ± SD of independent experiments. Statistical significance was determined using one-way ANOVA and Bonferroni’s post hoc analysis (a, b, c, d, h, i, m).

### WNK1 activation in senescent SARS-2-S syncytia impairs cardiac metabolism

Considering that cardiomyocytes with high energy demands are particularly susceptible to metabolism dysfunction and that the activation of WNK1 in the right ventricle (RV) causes RV dysfunction by promoting metabolic derangements^40, 41^, we next explored the possibility that senescent SARS-2-S syncytia in cardiomyocytes with WNK1 activation might contribute to impaired metabolism in the setting of heart dysfunction. Compared to nonfusion controls, the mitochondrial oxidative capacity of the cocultured cardiomyocytes was significantly decreased at 48 hpc as indicated by reduced basal respiration, ATP-linked respiration, spare respiratory, and maximal respiration (Fig. 6a, b). In addition, these senescent cells had more fragmented mitochondria with deteriorated glucose glycolysis capacity (Fig. 6c-f). Remarkably, small molecular inhibition of WNK1 via WNK463 was sufficient to rescue impaired mitochondrial respiration, mitochondrial integrity, and glycolytic metabolism (Fig. 6a-f).

**Fig. 6.**
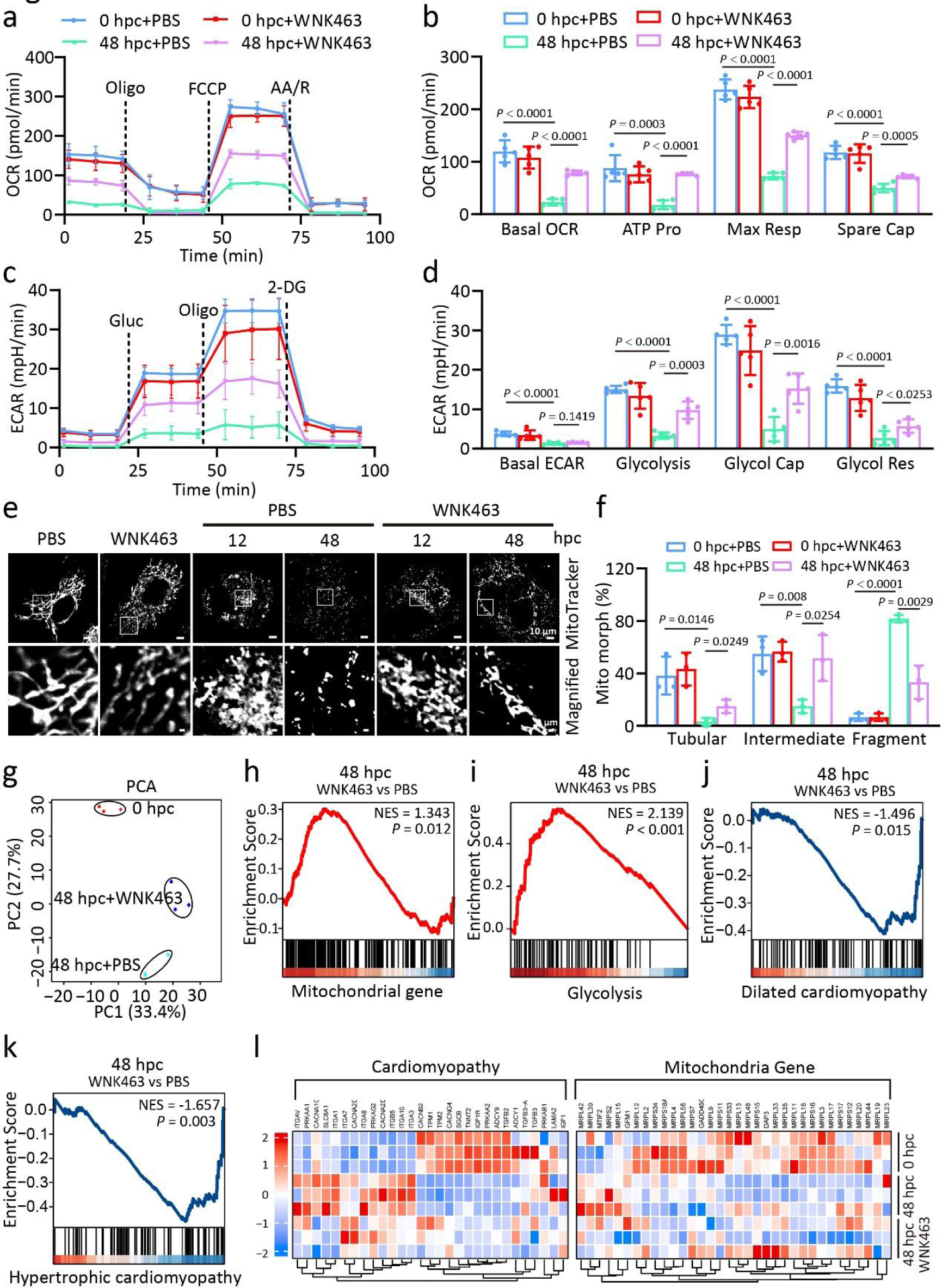
WNK1 activation in senescent SARS-2-S syncytia impairs cardiac metabolism. a-d, Seahorse analysis of OCR and ECAR in AC16 cocultured cells at 48 hpc in the presence or absence of WNK463 (1μM). OCR was measured with sequential addition of oligomycin (Oligo, 1.5μM), FCCP (2μM), and AA/R (1μM) (a) and the mitochondrial bioenergetic parameters including basal OCR, ATP production (ATP Pro), max respiration (Max Res), and spare capability (Spare Cap) were calculated (b). ECAR was measured with serial addition of addition of glucose (Gluc, 10 mM), oligo (1 μM), and 2-deoxyglucose (2-DG, 0.1 mM) (c) and the glycolytic parameters including basal ECAR, glycolysis, glycolysis capacity (Glyco Cap) and glycolytic reserve (Glyco Res) were calculated (d). e,f, Confocal microscopic images (e) and the quantification (f) of the mitochondrial morphology (Mito morph) in cocultured AC16 cells at 48 hpc in the presence or absence of WNK463 (1μM). Red, MitoTracker staining for mitochondria. Tubular: >10 μm length; Intermediate: ≤10 μm; Fragment: spherical (no clear length or width). Scale bars represent 10 μm (whole image) and 1 μm (magnified). g, PCA of gene expression patterns in cocultured AC16 cells at 48 hpc in the presence or absence of WNK463. In all plots, each point represented an independent biological replicate. h-k, GSEA of genes related to the mitochondrial (h), Glycolysis (i), dilated cardiomyopathy (j), and hypertrophic cardiomyopathy (k) signalling pathway among the DEGs in g. l, Heatmap of the critical DEGs in the cardiomyopathy and mitochondrial signalling pathway in cocultured AC16 cells at 48 hpc in the presence or absence of WNK463. All quantified data are shown as the mean ± SD of 3 independent experiments. All images are representative of 3 biological replicates. Statistical significance was determined using one-way ANOVA and Bonferroni’s post hoc analysis (b, d, e).

To further elucidate the mechanisms underlying the role of activated WNK1 in cardiac metabolism, we performed RNA-seq to profile the transcriptomes in syncytial cardiomyocytes with or without WNK463 challenge, using nonfusion cells as a control. PCA revealed that WNK463 shifted the transcript signature towards nonfusion controls (Fig. 6g). KEGG enrichment analysis showed the enrichment of hypertrophic cardiomyopathy and dilated cardiomyopathy in cocultured cardiomyocytes at 48 hpc (Extended Data Fig. 9). Remarkably, WNK463 treatment led to a significant enrichment of upregulated genes in mitochondrial integrity and glycolysis (Fig. 6h,i, l), whereas genes in cardiomyopathy were consistently enriched in downregulated DEGs (Fig. 6j-l). Taken together, we demonstrated that WNK1 activation in SARS-2-S syncytia significantly altered the heart transcriptomes and contributes to heart dysfunction by impairing cardiac metabolism.

### Senescent SARS-2-S syncytia exacerbated heart failure

To assess whether senescent SARS-2-S syncytium stimulates heart dysfunction*in vivo*, K18-hACE2 mice were administered with pseudovirus expressing BA.5 spike (SARS-2-Spp) intravenously (Fig. 7a). No significant differences were observed in LV mass or the heart-to-body weight (HW/BW) ratio between the PBS- and SARS-2-Spp-treated groups (Fig. 7b, c). HE and picrosirius red (PSR) staining of the myocardium showed that the extent of fibrosis was more profound in SARS-2-Spp-treated mice (Fig. 7d). Functional analysis of hearts, including ejection fraction (EF), fractional shortening (FS), left ventricular internal dimension at end diastole (LVID;d), left ventricular internal dimension at end-systole (LVID;s), as well as serum brain natriuretic peptide (BNP) levels between the two groups were also comparable (Fig. 7e-g and Extended Data Fig. 10a-g). Thus, SARS-2-S promotes fibrosis without causing baseline cardiac abnormalities during 7 observational days. We then repeated the above experiment with intraperitoneally (i.p.) injected isoproterenol (ISO) to assess the impact of SARS-2-Spp on cardiac function in mice with pre-existing heart failure. As expected, acute overstimulation with ISO significantly reduced the cardiac function and promoted the myocardial fibrosis (Fig. 7d-g). However, LV mass and HW/BW ratio were not affected (Fig. 7b, c). Strikingly, SARS-2-Spp treatment in ISO mice led to severe fibrosis and reduced heart function as compared to PBS-treatment in ISO mice (Fig. 7d-g). Notably, SARS-2-Spp also induced a significant increase in serum levels of troponin T (cTnT) and IL-1 β (Fig. 7h, i). Thus, SARS-2-Spp treatment exacerbated heart failure progression in pre-existing heart failure mice.

**Fig. 7.**
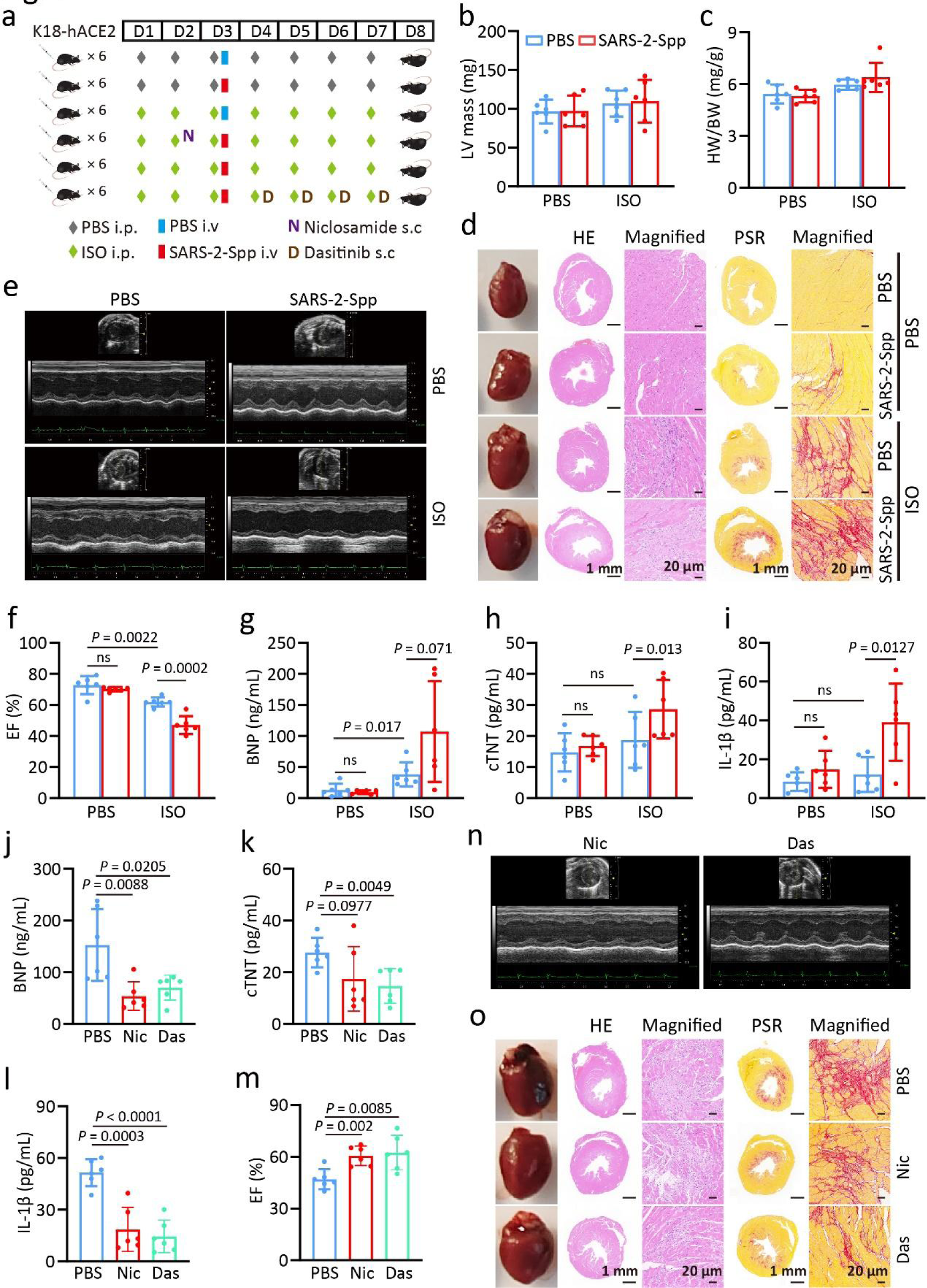
Senescent SARS-2-S syncytia exacerbated heart failure. a, Overview of the SARS-2-Spp-infected mouse model. ISO was subcutaneously injected into the backs of K18-hACE2 mice. Two days after ISO injection, 100 μL PBS or SARS-2-Spp (2.5X10^8^ TU/mL) was injected daily by tail vein daily continuing for 5 days. Niclosamide (100 mg/kg) was given by oral gavage 1 day before SARS-2-Spp injection, dasatinib (5 mg/kg) was given by oral gavage daily 1 day after SARS-2-Spp injection. Cardiac function was monitored via echocardiography 5 days after SARS-2-Spp injection. b-c, Left ventricular (LV) mass (b) and HW/BW ratio (c) in mice treated with PBS or SARS-2-Spp. d, Representative images showing whole hearts, cross-sections with HE staining and PSR staining from mice in b. Red, PSR staining for fibrotic areas. Scale bars, 1mm or 20 μm as indicated. e, Representative M-mode echocardiographic images from mice in b. f, Ejection fraction (EF) from mice in b. g-i, Serum BNP (g), cTnT (h), and IL-1β (i) levels from mice in b. j-l, Serum BNP (j), cTnT (k), and IL-1β (l) levels from from SARS-2-Spp-infected ISO-mice treated with PBS, niclosamide (Nic) or dasatinib (Das). m-n, EF (m) and representative M-mode echocardiographic images (n) from mice in j. o, Representative images showing whole hearts, cross-sections with HE staining and PSR staining from mice in j. Scale bars, 1mm or 20 μm as indicated. All quantified data are shown as the mean ± SD of n = 6 independent experiments. All images are representative of 3 biological replicates. Statistical significance was determined using one-way ANOVA and Bonferroni’s post hoc analysis (b, c, f, g, h, i, j, k, l, m).

To further prove that senescence SARS-CoV-2 syncytia act as a central pathogenic principle provoking heart failure, we attempted to eliminate senescent cells or inhibit fusion by using the senolytic drug dasatinib or the anti-syncytial drug niclosamide. We detected significantly lower concentrations of BNP, cTNT, and IL-1 β in the serum of mice treated with dasatinib and niclosamide (Fig. 7j-l). Intriguingly, the administration of dasatinib and niclosamide substantially alleviated the heart dysfunction and fibrosis severity in ISO-treated K18-ACE2 BALB/c mice challenged with SARS-2-Spp (Fig. 7m-o and Extended Data Fig. 10h-n), further supporting the notion that senescent SARS-2-S syncytia contribute to exacerbated heart failure.

### WNK1 inhibition effectively protects the heart from exacerbated heart failure triggered by SARS-2-S

We then wanted to test if WNK1 inhibition by WNK463 can alleviate heart failure exacerbated by SARS-2-S through rescuing metabolic dysfunction. Strikingly, mice treated with WNK463 showed improved heart function as determined by enhanced EF, FS, and reduced serum BNP levels (Fig. 8a-e). WNK463 exposure also led to significantly reduced serum levels of cTNT and IL-1 β (Fig. 8f, g). Therefore, WNK1 intervention can effectively protect the heart from exacerbated heart failure triggered by SARS-2-S.

**Fig. 8.**
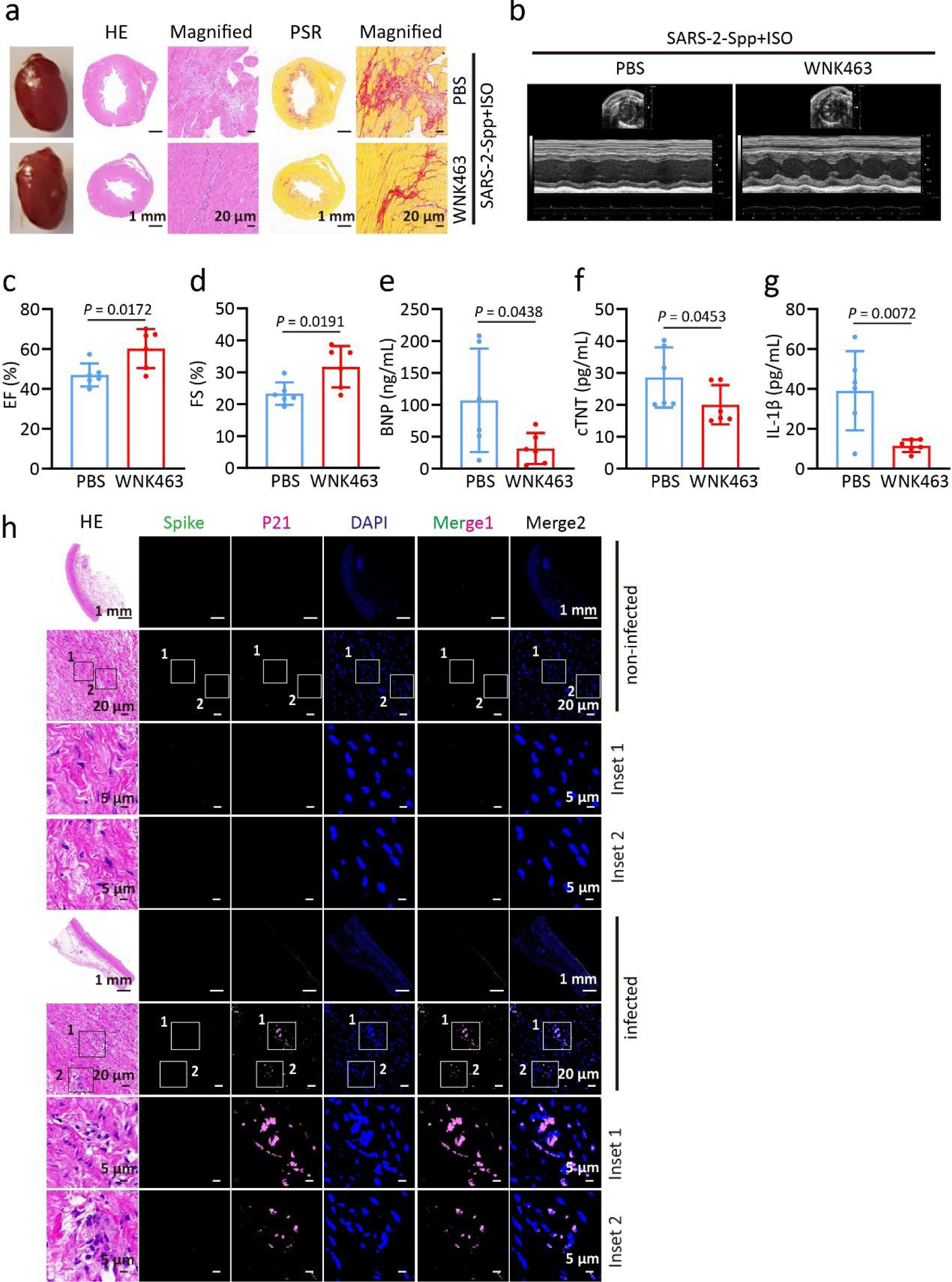
WNK1 inhibition effectively protects the heart from exacerbated heart failure triggered by SARS-2-S. a, Representative images showing cross-sections with HE staining and PSR staining from SARS-2-Spp-infected ISO mice treated with PBS or WNK463. b-l, Representative M-mode echocardiographic images (b), EF (c), FS (d), serum BNP (e), cTnT (f), and IL-1β (g) levels from mice in a. h, HE staining and immunofluorescence analysis of ascending aorta tissue from SARS-CoV-2-infected and non-infected patients. Green, Spike; pink, p21; blue, DAPI staining. Scale bars represent 1 mm or 20 μm as indicated. All images from mouse tissue are representative of 3 biological replicates. All quantified data are shown as the mean ± SD of n = 6 independent experiments. Statistical significance was determined using two-tailed Student’s t test (c, d, e, f, g, h, I, j, k, l).

### Syncytia in SARS-CoV-2-infected patients were senescent

We next asked whether senescent syncytia would be a typical feature detectable in SARS-CoV-2-infected patients. We specifically analysed ascending aorta tissue specimens from one SARS-CoV-2-infected patient and one non-infected patient with hypertension undergoing aortic dissection surgery. Strikingly, typical syncytia cells with a large cytoplasm containing a variable number of nuclei (2 to more than 20) were observed in ascending aorta tissue from infected individuals without remaining SARS-2-S expression, but not in non-SARS-CoV-2-infected patient with the same age and gender. Notably, these syncytia cells exhibited strong p21 staining intensity (Fig. 8h). Collectively, our data constitute the first*in vivo* evidence showing the existence of syncytia with a senescence-like phenotype in SARS-CoV-2-infected patients with cardiovascular disease.

## Discussion

COVID-19 patients with pre-existing heart failure are at higher risk of developing severe symptoms and experiencing mortality compared to COVID-19 patients without heart failure history^42^. Direct myocardial injury or inflammatory-mediated injury has been suggested to explain the increased vulnerability to heart failure in patients with COVID-19^43^. Here, we directly linked SARS-2-S-triggered syncytium formation with the ensuing induction of cellular senescence and its pathophysiological contribution to heart failure progression. We found that both SARS-2-S expression and SARS-2-S protein internalization were sufficient to induce senescence in nonsenescent ACE2-expressing cells. In particular, mRNAs-SARS-2-S injection provoked pathways enriched in senescence and SASP phenomenon*invivo*. Mechanistically, MAVS regulated the anti-death to senescence fate of SARS-2-S syncytia through the TNFα-TNFR2 axis. SARS-2-S senescent syncytia with a smaller size tendency triggered WNK1 activation and led to impaired cardiac metabolism. Evidences of senescent syncytia puncta were observed in ascending aorta specimens from patients infected with SARS-CoV-2 omicron variant. Senescence induction by SARS-2-S depends on its fusogenic ability as the SARS-2-SRA mutant was unable to induce fusion and failed to induce senescence and WNK1 activation. Notably, pre-existing heart failure was significantly exaggerated by SARS-2-Spp in K18-hACE2 mice.*Invivo*testing of senolytic, WNK1 inhibitor, and anti-syncytia drug implicated senescence syncytia as a central driver in promoting heart failure progression, highlighting the clinical potential of WNK1 inhibitors as a novel interventional therapeutic strategy.

Activation of DNA sensors by DNA chromatin has been illustrated in promoting paracrine senescence and the production of SASP in VIS^44^. Here, we show that RIG-I-MAVS signalling was activated and governed the senescence fate of SARS-2-S syncytia. While we cannot definitively exclude the possible involvement of chromatin DNA and its sensors cGAS and STING in activating the senescence response induced by SARS-2-S syncytia, the*invitro* knockdown experiment clearly demonstrates the crucial role of RIG-I-MAVS, but not STING, in this process. The existence and sources of endogenous dsRNA warrant further investigation. Nevertheless, we provide evidence of the contribution of RIG-I-MAVS to the regulation of SARS-2-S syncytium fate through TNFα. Using multiplex protein analysis, we identified a burst of SASP-related cytokines, including TNFα, IL-6, IFN-γ, and others, in the hACE2 mouse model injected with SARS-2-S mRNA. Although the exact contribution of each cytokine to SARS-2-S syncytium survival and senescence remains unclear, we provide evidence that the central regulation of SARS-2-S syncytium survival and senescence may be dependent on TNFα. Previous studies reported that TNFR2 interacts with the transmembrane form of TNFα (mTNFα) upon cell-to-cell contact^45–47^. Indeed, TNFR2 siRNA effectively reduced the increased survival of cocultured cells induced by TNFα, highlighting the indispensable role of TNFα/mTNFα in governing the fate of SARS-2-S syncytia mainly by TNFR2.

With the progression of COVID-19, heart failure appears to be a potential hazard due to SARS-CoV-2 infection, chiefly in elderly patients with hypertension, ischaemic heart disease, and diabetes mellitus^48^. Patients with basic heart failure disease showed elevated ACE2 expression^49^, whereas patients with PACS exhibited persistent circulating SARS-2-S and EVs harbouring SARS-2-S^16, 17^. High ACE2 expression and long SARS-2-S duration may increase the likelihood of syncytium formation. Moreover, the pre-existing inflammatory environment with older age or with chronic disease, especially those with high TNFα levels, may increase the survival and senescence of syncytia. We therefore proposed that factors including pre-existing inflammation, high ACE2 expression, and persistent SARS-2-S could contribute to cardiac or multi-organ damage during different stages of SARS-CoV-2 infection by increasing the burden of senescent syncytia. In support of this, our study provide the first *invivo* evidence showing the existence of senescent syncytia, albeit unknown cell types, in aorta arch tissue of one female patient with hypertension during SARS-CoV-2 infection. In addition, vaccines using recombinant SARS-2-S fragments or other derivatives of SARS-2-S that are not fusogenic were reported to have fewer complications than vaccines that express fully or partially fusogenic SARS-2-S. S It remains to be demonstrated whether the senescent syncytia were induced in cell autonomous manner or through paracrine effect and its pathological consequences observed in assay systems are of relevance for clinical heart failure progression.

A defining feature of cellular senescence is characterized by its distinct morphological changes during progression^38^. Here, we found that SARS-2-S senescent syncytia exhibited a smaller surface area per nucleus. As a molecular crowding sensor that undergoes cell shrinkage-dependent phase separation to restore cell volume, WNK1 phase separation was identified in SARS-2-S syncytia. It has also been reported that TNFα activates the WNK1-SPAK-NCC phosphorylation cascade and contributes to salt-sensitive hypertension in several CKD mouse models^50^. Therefore, the effect of TNFα on WNK1 activation should be carefully monitored later. Previous studies have shown that small-molecule inhibition of WNK1 prevents the downregulation of mitochondrial metabolism and improves RV function and survival in pulmonary arterial hypertension^41^. We found that WNK1 inhibition alleviated SARS-2-S-aggravated heart failure and reduced serum cTnT and IL-1β levels. cTnT is released into circulation during injury to cardiac myocytes^51^. Individuals with higher cTnT levels are at an increased risk of developing heart failure and sudden cardiac death ^52^. Therefore, through its metabolism-rescue and anti-inflammatory properties, WNK463 may have dual protective effects in preventing severe myocardial lesions and heart failure in “high-risk” patients. As TNFα and anti-syncytial drug acts in the early phase of cell-cell fusion, its narrow time window may limit its therapeutic use. While senescence elimination reduces hypertrophy and improves function, how the heart responds to senolytic treatment in longer-term studies has yet to be investigated. Here, we recommend that rescue of cardiac metabolism dysfunction should be taken into consideration in individuals during the acute or post-acute stage of SARS-CoV-2 infection. It should be noted that SARS-CoV-2 infection also induces new onset of multiple cardiovascular diseases and causes long-term cardiovascular sequelae in PACS patients^53^. Cytokine storm was suggested to be a point of convergence in the pathophysiologic process between COVID-19 and heart failure^54^. Here we found that acute SARS-2-Spp treatment led to profound cardiac fibrosis without affecting baseline cardiac function. Further studies examining the chronic pathological role of SARS-2-S senescent syncytia are needed to determine whether WNK inhibitors are beneficial in preventing the new onset of cardiovascular diseases or alleviating long-term cardiovascular sequelae in PACS patients.

In summary, we provide evidence showing direct induction of cell-cell fusion with a senescence-like phenotype in FFWI and FFWO manners by SARS-2-S both*in vitro* and *in vivo*. The susceptibility of patients with heart failure to more pronounced complications during the acute and post-acute stages of SARS-CoV-2 infection could be related to the senescent outcome of SARS-2-S syncytia. WNK1 may be a druggable target in the management of heart failure associated with SARS-CoV-2 infection.

## Materials and Methods

### Reagents

Dulbecco’s modified Eagle’s medium (DMEM, high glucose, D5796) and protease inhibitor cocktail I (20-201) were purchased from Millipore (Massachusetts, USA). Foetal bovine serum (FBS, A3160901) was purchased from Gibco (New York, USA). Lipofectamine™ 2000 (11668-027), MitoTracker^TM^ Deep Red FM and CellEvent™ Senescence Green detection kits (C10850) were purchased from Invitrogen (Massachusetts, USA). BCA protein concentration determination kit (P0012), DAPI (C1002) and β-galactosidase reporter gene assay kit (RG0036) were purchased from Beyotime (Shanghai, China). NucleoZOL (740404) was purchased from MACHEREY-NAGEL (MN, Düren, Deutschland). PerfectStart^TM^ Green qPCR SuperMix (AQ601) and TransScript^®^ One-Step gDNA Removal and cDNA Synthesis SuperMix (AT311) were purchased from TransGen Biotech (Beijing, China). 4-PBA (HY-A0281), WNK463 (S8358), fisetin (S2298), dasatinib (S1021) and venetoclax (S8048) were purchased from Selleck Chemicals (Houston, TX). Recombinant human IL-6 (AF-200-06) and human TNFα (300-01A) were purchased from PeproTech (New Jersey, USA). Mouse cTnT/TNNT2 ELISA Kit (E-EL-M1801c), and Mouse BNP ELISA Kit (E-EL-M0204c) were purchased from Elabscience (Wuhan, China). Mouse 1L-1 β Precoated ELISA Kit (1210122) was purchased from Dakewe Biotech (Beijing, China).

### Antibodies

Anti-CD81 (ab109201, 1:1,000 dilution), anti-CD63 (ab134045, 1:1,000 dilution), anti-TSG101 (ab125011, 1:1,000 dilution), anti-CD9 (ab263019, 1:1,000 dilution), anti-ACE2 (ab108252, 1:1,000 dilution), anti-WNK1 (ab174854, 1:2,000 dilution), anti-SPAK (ab128894, 1:2,000 dilution) and anti-IRF3 (ab76409, 1:1000 dilution) were purchased from Abcam (Cambridge, UK). Anti-p-SPAK ser373, anti-α-tubulin (T9026, 1:5,000 dilution) and anti-Flag (A8592, 1:10,000 dilution) were purchased from Sigma-Aldrich (Missouri, USA). Anti-TNFα (60291-1-Ig, 1:5,000 dilution), anti-MAVS (14341-1-AP, 1:100 dilution for immunofluorescence, 1:4,000 dilution for immunoblotting), anti-TRAF3 (18099-1-AP, 1:1,000 dilution), anti-p21 (10355-1-AP, 1:100 dilution), anti-p16 (10883-1-AP, 1:100 dilution), anti-TBK1 (28397-1-AP, 1:2,000 dilution), anti-STING (19851-1-AP, 1:3,000 dilution), CoraLite^®^488-conjugated TNFα antibody (CL488-60291, 1:200 dilution), anti-TNFR2 (28746-1-AP, 1:200 dilution) and anti-GAPDH (60004-4-I-Ig, 1:5,000 dilution) were purchased from Proteintech (Wuhan, China). Anti-SARS-CoV/SARS-2-S (GTX632604, 1:2,000 dilution) was purchased from GeneTex (CA, USA). Anti-TRAF6 (8028S, 1:1,000 dilution), anti-phospho-p65 (3033T, 1:1,000 dilution), anti-RIG-I (3743T, 1:1,000 dilution), anti-phospho-IRF3 (4947S, 1:1,000 dilution) and anti-phospho-TBK1 (5483T, 1:2,000 dilution) were purchased from Cell Signaling Technology (Massachusetts, USA). Anti-phospho-p65 (A19653, 1:1,000 dilution) was purchased from ABclonal (Wuhan, China). Anti-rabbit HRP-IgG (ZB-2301, 1:5,000 dilution) and anti-mouse HRP-IgG (ZB-2305, 1:5,000 dilution) were purchased from ZSGB-BIO (Beijing, China).

### Plasmids, cell culture and small interfering RNAs (siRNAs)

Mammalian expression vectors encoding SARS-2-S (Wuhan-Hu-1, GenBank: NC_045512.2), SARS-2-S_NF_ (spike R682A, R683A, R685A, K814A and R815A) and SARS-2-S_O_ (Omicron) were codon-optimized and cloned into the pcDNA3.1 expression vector. To produce lentivirus plasmids, SARS-2-S and ACE2 were cloned into the pCDH-CMV-MCS-EF1-Puro vector. NF-κB-Luc, IFN-β-Luc, and IRF-3-Luc reporter gene constructs were kept in our laboratory. All plasmids were verified by DNA sequencing.

The A549 (CRL-185), HEK293T (293T, CRL-3216), HeLa (CRL-3216), and HIEC-6 (CRL-3266) cell lines were obtained from the American Type Culture Collection (ATCC, Rockville, MD, USA). The AC16 (SCC109) cell line was obtained from Sigma-Aldrich (Missouri, USA). MAVS-KO 293T cells were kindly provided by Prof. T. Li (National Center of Biomedical Analysis, Beijing, China). All cell lines were mycoplasma free and incubated in DMEM supplemented with 10% FBS, 100 U/mL penicillin, and 0.1 mg/mL streptomycin at 37°C under a humidified atmosphere with 5% CO2. Lipofectamine^TM^ 2000 was used for transfection following the manufacturer’s protocol. For lentivirus production, pMD2. G, psPAX2 and pCDH-SARS-2-S or pCDH-ACE2 were transfected into 293T cells together, and lentiviral particles were harvested from cell supernatants filtered through a 0.45 μm filter 48 h later. For stable overexpression of ACE2 and SARS-2-S, 293T, A549, AC16, HIEC-6, and HeLa cells were infected with harvested lentiviral particles and selected in the presence of puromycin (2-4 μg/mL). Stably transfected clones were selected under puromycin pressure after two weeks. To knock down specific genes, siRNAs targeting human genes were purchased from GenePharma, and their target sequences are listed in Table S1.

### Cell-cell fusion assay

Cells were stably transfected with SARS-2-S or ACE2 and cultured to 60% confluence. Then, SARS-2-S-expressing cells and ACE2-expressing cells were either mixed at 1:1 ratios or allowed to continue culture. At the indicated times, the cocultured cells were harvested, and the individually cultured SARS-2-S-expressing cells and ACE2-expressing cells were mixed*invitro*using the same ratio used as the nonfusion controls.

### Quantification of fusion level based on β-galactosidase reporter gene assay

To quantify the cell-cell fusion level, an α-complementation of the lacZ system based on β-galactosidase was used^14^. Producer cells were constructed with SARS-2-S and lacZ (△ 11-41 aa), and target cells were constructed with ACE2 and lacZ (1-56 aa). Upon cell-cell fusion, complementation of lacZ (△11-41 aa) with lacZ (1-56 aa) inside the syncytium results in the formation of active β-galactosidase, which is then quantified by a β-galactosidase Reporter Gene Assay Kit according to the manufacturer’s instructions. Briefly, 50 μL of lysate collected from the syncytium at the indicated times was treated with equal amounts of the specific substrate 2-nitrophenyl β-D-galactopyranoside. After 3 h of incubation, the reactions were quenched with 150 μL stop solution, and the absorbance was measured at 420 nm using a multimode microplate reader (Tecan).

### SARS-2-S-mRNA-LNP synthesis

T7 RNA polymerase-mediated transcription from a linearized DNA template from a plasmid that encodes codon-optimized SARS-2-S was used to produce mRNA *invitro*. LNP formulations were prepared using a method previously described for siRNA with modifications^55^. Briefly, lipids were dissolved in ethanol containing an ionizable lipid, 1,2-distearoyl-sn-glycero3-phosphocholine, cholesterol and PEG-lipid (at molar ratios of 50:10:38.5:1.5). The lipid mixture was combined with 20 mM citrate buffer (pH 4.0) containing mRNA at a ratio of 1:2 through a T-mixer. The formulations were then diafiltrated against a 10 × volume of PBS (pH 7.4) through a tangential-flow filtration membrane with a 100 kDa molecular weight cut-off (Sartorius Stedim Biotech), concentrated to the desired concentrations, and passed through a 0.22 μm filter. All formulations were tested for particle size, distribution, RNA concentration and encapsulation.

### EV purification

For EV production, supernatants of HeLa and SARS-2-S stably transfected HeLa cells cultured in EV-depleted medium were collected after 72 h and centrifuged at 300 × g for 10 min to remove the cells. The cell supernatant was diluted 1:2 with cold PBS before low-speed centrifugation at 3000 × g for 10 min, and the resulting supernatant was transferred to new tubes for ultracentrifugation at 100,000 ×g using a type 100 TI rotor in an XPN-100 instrument (CP100NX; Hitachi, Brea, USA) for 90 min. The resulting EV pellet was resuspended in 250 mL of cold PBS and denoted EVs.

### Flow cytometry

To assess the cellular uptake of S-EVs, the cells were exposed to DiO-labelled EVs at the indicated doses and then incubated at 37°C for the indicated durations. After extensive washing, flow cytometry was performed with an Attune^®^ NxT instrument (Thermo Fisher Scientific, Massachusetts, USA), and 30,000 cell acquisitions under a medium flow rate were used for data capture. Data analysis was performed using FlowJo software (V10.0.7r2). A control sample consisting of DiO dye used as the background was prepared following the same procedure indicated above.

For SARS-2-S protein expression detection, cells were treated with S-EVs. Forty hours later, the cells were detached and incubated with SARS-2-S mAb (GTX632604, clone 1A9, 1:1,000 dilution) for 30 min on ice, followed by incubation with PE goat anti-mouse IgG (Biolegend, 405307, 1:100 dilution) for 15 min at room temperature. The cells were then analysed by flow cytometry.

### Multiplex bead-based protein detection (MAGPIX)

Multiplex bead-based protein analyses were conducted according to the manufacturer’s instructions with a custom Procartaplex 36-plex panel (Thermo Fisher Scientific, cat no. EPX360-26092-901). Briefly, samples were diluted, and 50 µL of each prepared sample was incubated with antibody-conjugated magnetic beads for 1 h overnight at 4°C. After washing, detection antibodies were added and incubated at room temperature for 30 min. Following further washing steps, PE-streptavidin was added and incubated for 30 min. The assay data were measured with the Bio-Plex^®^ MAGPIX™ multiplex reader (Bio-Rad) and analysed with a 5PL algorithm provided by Bio-Plex Manager™ software (version 6.1).

### Semidenaturing detergent agarose gel electrophoresis (SDD-AGE)

Crude mitochondria and cytosolic extracts were separated from cells using differential centrifugation with a cell mitochondria isolation kit (Beyotime, C3601) according to a published protocol^56^. A549 cells were resuspended and lysed by grinding and were centrifuged at 800 × g for 10 min at 4°C. The supernatants were transferred to a new tube and centrifuged at 13,000 × g for 10 min at 4°C to separate the crude mitochondria and cytosolic extracts. The crude mitochondria were resuspended in sample buffer (0.5 × TAE, 10% glycerol, 2% SDS, and 0.0025% bromophenol blue) and loaded onto a 1.5% agarose gel. Then, the samples were electrophoresed at a constant voltage of 80 V at 4°C. The proteins were transferred to PVDF membranes for immunoblotting.

### Luciferase reporter assays

A luciferase expression vector controlled by a promoter containing three repeats of the NF-κB response element was obtained from Beyotime (D2204). The pSV40-Renilla plasmid was purchased from Promega (E2231). To assess the activity of NF-κB, cells were plated in triplicate in a 24-well plate and then transiently transfected with pNF-κB-luc and the internal control plasmid pSV40-Renilla for 12 h. The cells were then cocultured for the indicated durations.

The cells were harvested, and firefly luciferase activity and Renilla luciferase activity were detected using the dual-luciferase reporter assay system (Promega, E1910) with a multimode microplate reader (Tecan). The firefly luciferase luminescence data were normalized to the Renilla luciferase luminescence data.

### VSV infection

Cells were plated in 12-well plates and VSV was added to the medium at a concentration of 80 haemagglutinating units per mL and a multiplicity of infection (MOI) of 5 at 60% cell confluence. After 1 h of incubation, the extracellular virus was removed by washing the cells twice with serum-containing medium. Cells were fixed at the indicated times after infection for immunofluorescence assays.

### Syncytial viability assay

A CCK8 assay was used to assess syncytial cell viability. Briefly, cocultured cells were seeded into a 96-well plate at a cell density of 3000 per well and allowed to adhere for 72 h. Then, the medium was washed and replaced with fresh medium, followed by treatment with serially diluted inhibitors. At 24 h after treatment, 10 µL of CCK8 solution was added to 100 µL of culture medium, and the absorbance of each well was measured at 450 nm using a multimode microplate reader (Tecan) with Tecan Spark Control (v.2.1) software. All experiments were performed in triplicate, and the cell viability was calculated as the ratio of the viability under each experimental condition to the viability of the control.

### Animals

Male C57BL/6J WT mice (8-week-old) were purchased from SPF Biotechnology (Beijing, China). All mice were conventionally group-housed on a 12-h light/dark cycle in an animal facility for 3 days before any procedures.

For S-mRNA-LNP injections, we injected 3.0 x 10^11^ μg AAV-hACE2 constructs into the tail veins of 8-week-old C57BL/6J mice to drive mainly hepatic hACE2 expression. Four weeks later, the mice were administered 3.0 μg of S-mRNA-LNP or LNP via i.m. injection, followed by terminal sacrifice of the mice and the collection of sera and livers at the indicated times.

For ISO-induced heart failure model, 12-week-old K18-hACE2 C57BL/6J male mice was subcutaneously injected with ISO (5 mg/kg) into the backs daily continuing for 7 days. Two days after ISO injection, the mice were injected with 100 μL PBS or SARS-2-Spp (2.5X10^8^ TU/mL) by tail vein for 5 days. For drug testing, niclosamide (100 mg/kg) was given by oral gavage 1 day before SARS-2-Spp injection. Dasatinib (5 mg/kg) was given by oral gavage daily 1 day after SARS-2-Spp injection. WNK463 (2.5 mg/kg) was given intraperitoneally daily 1 day after SARS-2-Spp injection. Five days after SARS-2-Spp injection, the mice were anaesthetized with isoflurane and cardiac function was analyzed by a Vevo2100 digital high-frequency ultrasound system (FUJIFILM VisualSonics, Inc). Mice were then sacrificed, serum and the organs were collected for future use. All animal studies were performed at the AMMS Animal Centre according to standard operating procedures in a specific pathogen-free facility under the approval of the Institutional Animal Care and Use Committee.

### Quantitative real-time PCR (RT-qPCR)

Total mRNA was extracted from cells or tissues using NucleoZOL. cDNA was prepared from total mRNA by using TransScript^®^ One-Step gDNA Removal and cDNA Synthesis SuperMix, and the relative amounts of individual mRNAs were calculated after normalization to the corresponding β-actin mRNA. The primer sequences are included in Table S2.

### Western blotting and immunoprecipitation

Cells were lysed in NP40 cell lysis buffer with fresh protease inhibitors. Supernatants were separated by SDS-PAGE after centrifugation and transferred to PVDF membranes for immunoblot analyses using the indicated antibodies. Whole-cell lysates for coimmunoprecipitation were prepared by sonication in NP40 buffer (1% NP40, 150 mM NaCl and 40 mM Tris pH 7.5). Clarified lysates were then incubated with the indicated antibodies and with Protein A/G PLUS-agarose. After four washes with PBS, the immunoprecipitates were collected by centrifugation and reserved for immunoblot analyses.

### RNA-sequencing (RNA-seq) analysis

Total RNA was isolated from cells or liver tissues with TRIzol reagent, and the RIN was checked to inspect RNA integrity with an Agilent 2100 Bioanalyzer (Agilent Technologies, Santa Clara, CA, US). The qualified total RNA was further purified with an RNAClean XP kit (cat No. A63987, Beckman Coulter, Inc., Kraemer Boulevard, Brea, CA, USA) and RNase-Free DNase Set (cat# 79254, QIAGEN, GmBH, Germany). cDNA libraries were constructed following Illumina standard protocols and sequenced with an Illumina NovaSeq 6000 by SHBIO (Shanghai, China). The sequencing reads were mapped to the human genome using TopHat (version 1.0.13). Avadis NGS (version 1.3) was used to calculate reads per kilobase per million mapped reads (RPKM) values. The P value and fold change (FC) in the expression of genes between groups were jointly used to identify differentially expressed genes (DEGs). The criteria for statistical significance were P< 0.05 and |Log2(FC)|>1. GO enrichment and KEGG pathway analyses were conducted by SHBIO to investigate the potential functions of the DEGs, and P< 0.05 was considered to indicate a statistically significant difference.

### Immunofluorescence analysis

For immunofluorescence staining, cells or tissue sections were fixed with a 4% paraformaldehyde solution, washed twice with PBS and permeabilized with 0.3% Triton X-100 in PBS. The Triton X-100 solution was replaced with 5% (w/v) BSA, and samples were incubated with primary antibodies for 1 h. After washing three times with PBST (0.05% Tween 20 in PBS), the samples were incubated with fluorophore-conjugated secondary antibodies diluted in PBS for 30 min. After washing three times with PBST, the nuclei were labelled with a DAPI solution. For dsRNA staining assays, J2 was used as the primary antibody, and Alexa Fluor™ 555-labelled donkey anti-mouse IgG (H+L) secondary antibody was used as the secondary antibody.

For SA-β-gal staining assays, SA-β-gal was detected with the CellEvent™ Senescence Green Probe under a fluorescence microscope. Fluorescence images were acquired under a Nikon A1 confocal microscope, Zeiss LSM710 confocal microscope or Tecan Spark Cyto. The images were processed and analysed using Volocity (v.6.1.1, PerkinElmer), Nikon NIS-Elements AR (v4.00.12) or ImageJ (version 1.53c).

### Seahorse assay

The Seahorse XFe24 Extracellular Flux Analyzer and Seahorse XF Glycolytic Rate Assay Kit from Agilent Technologies were used to measure ECAR and OCR according to the manufacturer’s instructions. In brief, cells were seeded in XFe24 microplates and incubated overnight at 37°C in a CO2 incubator. On the day of the assay, the cells were washed and incubated with Seahorse assay medium. OCR was measured in response to oligomycin (1.5 μM), FCCP (2 μM), and antimycin A /rotenone (AA/R, 1 μM), while ECAR was measured in response to glucose (10 mM), oligomycin (1 μM), and 2-deoxyglucose (2-DG, 0.1 mM). The results were analysed using Wave software (Seahorse/Agilent).

### Human aortic tissue samples acquisition

Aortic tissue samples were obtained from 2 patients of the same age and gender undergoing aortic dissection surgery with or without SARS-CoV-2 infection. Basic informations were collected from electronic medical files of the department of pathology at FuWai Hospital (Table S3). Written informed consent was provided and all experiments conducted with human tissue samples were performed in accordance with the relevant guidelines and regulations.

### Statistical analysis

In the present study, GraphPad Prism 8.0 was used for statistical calculations and data plotting. Differences between two independent samples were evaluated by two-tailed Student’s t tests or the Mann-Whitney test, as appropriate. Differences between multiple samples were analysed by one-way ANOVA or two-way ANOVA, followed by Bonferroni’s post hoc analysis, as appropriate. All tests were two-tailed unless otherwise indicated. We considered a P value<0.05 to indicate statistical significance. Significance values were set as follows: ns (not significant), P > 0.05; *, P < 0.05; **, P < 0.01; ***, P < 0.001.

Competing Interest Statement: The authors declare no competing interests.

---

**Fig. S1.**
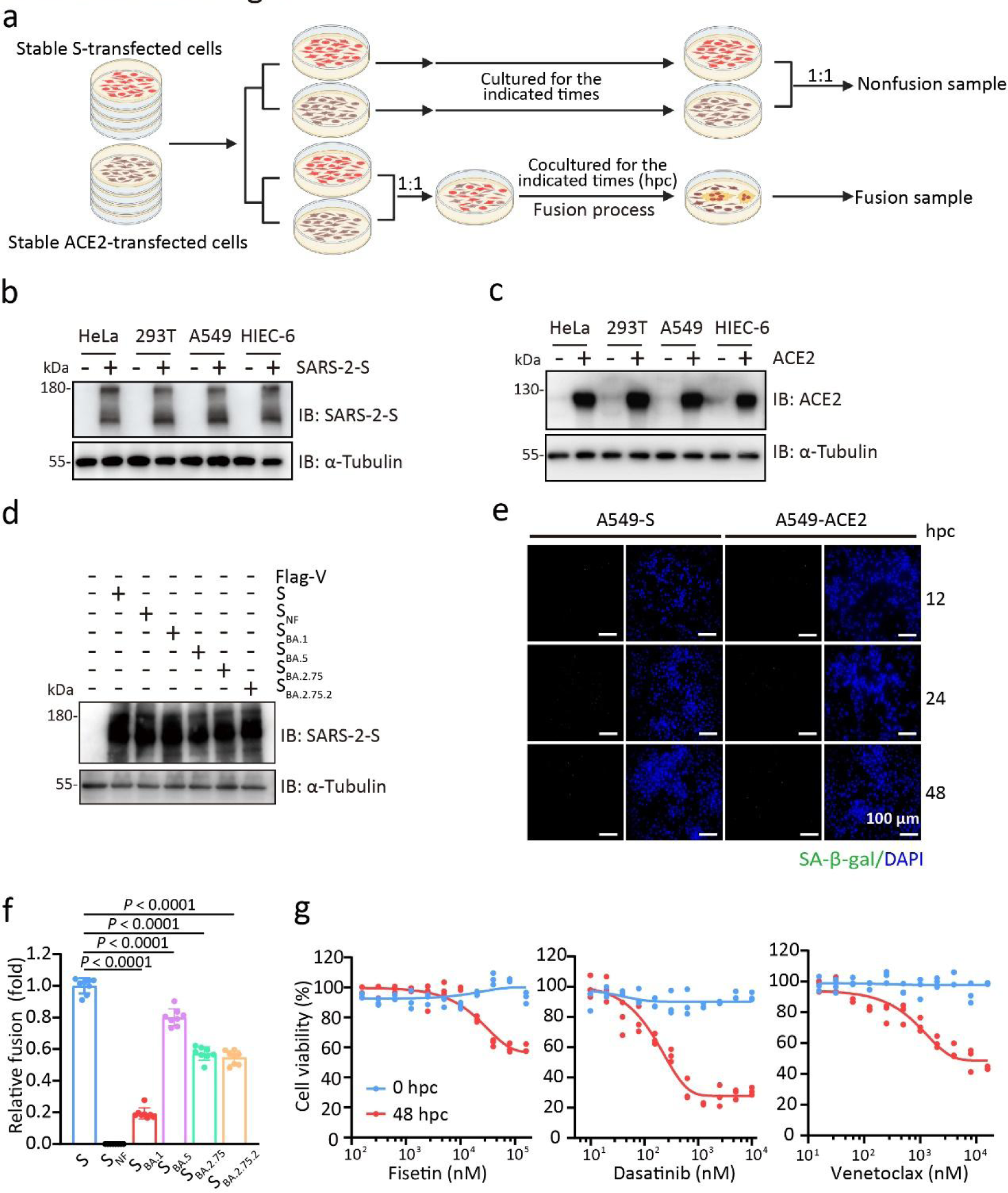
SARS-2-S syncytia exhibit a senescence-like phenotype. a, Illustration of the cell coculture system. Cells were stably transfected with plasmid DNA expressing S or human ACE2, followed by coculture at a 1:1 ratio for the indicated times. A mixture of A549-S cells and A549-ACE2 cells mixed at a 1:1 ratio without coculture (0 hpc) was used as a nonfusion control. b-c, Immunoblot analysis of S (b) and ACE2 (c) expression in A549, 293T, HeLa, and HIEC-6 cells transfected with the indicated plasmids. α-Tubulin was used as a loading control. d, Immunoblot analysis of A549 cells transfected with S, S_NF_, S_BA.1_, S_BA.5_, S_BA.2.75_ or S_BA2.75.2_. α-Tubulin was used as a loading control. e, SA-β-gal staining of A549-S or A549-ACE2 cells. Green, SA-β-gal staining; blue, nuclear DAPI staining. Scale bars represent 100 μ m. f, Quantification of the relative fusion ability of four Omicron variants based on the β-galactosidase α-complementation assay. The fusion level of A549-S and A549-ACE2 cocultured cells at 24 hpc was set to 1. g, Cell viability of cocultured A549 cells at 48 hpc treated with fisetin, dasatinib, or venetoclax for 12 h by CCK8 assay. All images are representative of 3 biological replicates. All quantified data are presented as the mean ± SD of n = 3 independent experiments. Statistical significance was determined with two-tailed Student’s t test (f).

**Fig. S2.**
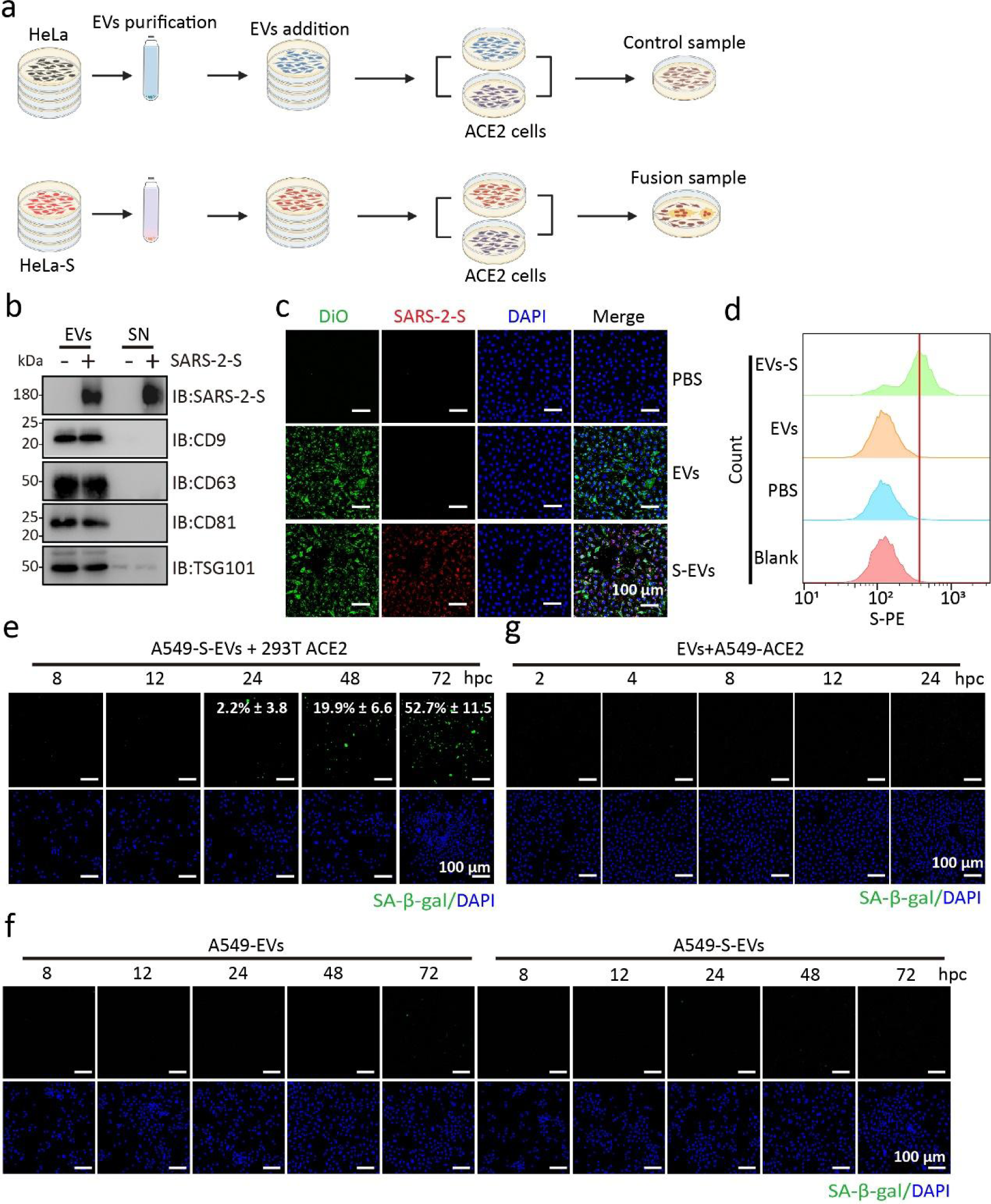
SARS-2-S delivered by EVs triggers syncytial senescence. a, Illustration of EV-mediated FFWO. Cells were treated with S-EVs for 24 h, followed by coculture with ACE2-expressing cells for the indicated times. A mixture of A549-EVs and A549-ACE2 cells was used as a nonfusion control. b, Immunoblot analysis of EV markers (CD9, CD63, CD81, and TSG101) and S expression in EVs purified from the supernatant (SN) of HeLa or HeLa-S cells. c, Analysis of DiO-labelled S-EVs or EVs uptake by A549 cells. Green, DiO-labelled EVs; red, S staining; blue, nuclear DAPI staining. d, Flow cytometry analysis of S expression on the cell surface of A549-S-EVs cells. e, g, SA-β-gal staining of A549-S-EVs cocultured with 293T-ACE2 cells (e) or EVs-treated A549-ACE2 cells (g) for the indicated times. Green, SA-β-gal staining; blue, nuclei DAPI staining. Scale bars represent 100 μm. f, SA-β-gal staining of EVs- or S-EVs-treated A549 cells for the indicated times. Green, SA-β-gal staining; blue, nuclei DAPI staining. Scale bars represent 100 μ m. All the images are representative of three independent experiments.

**Fig. S3.**
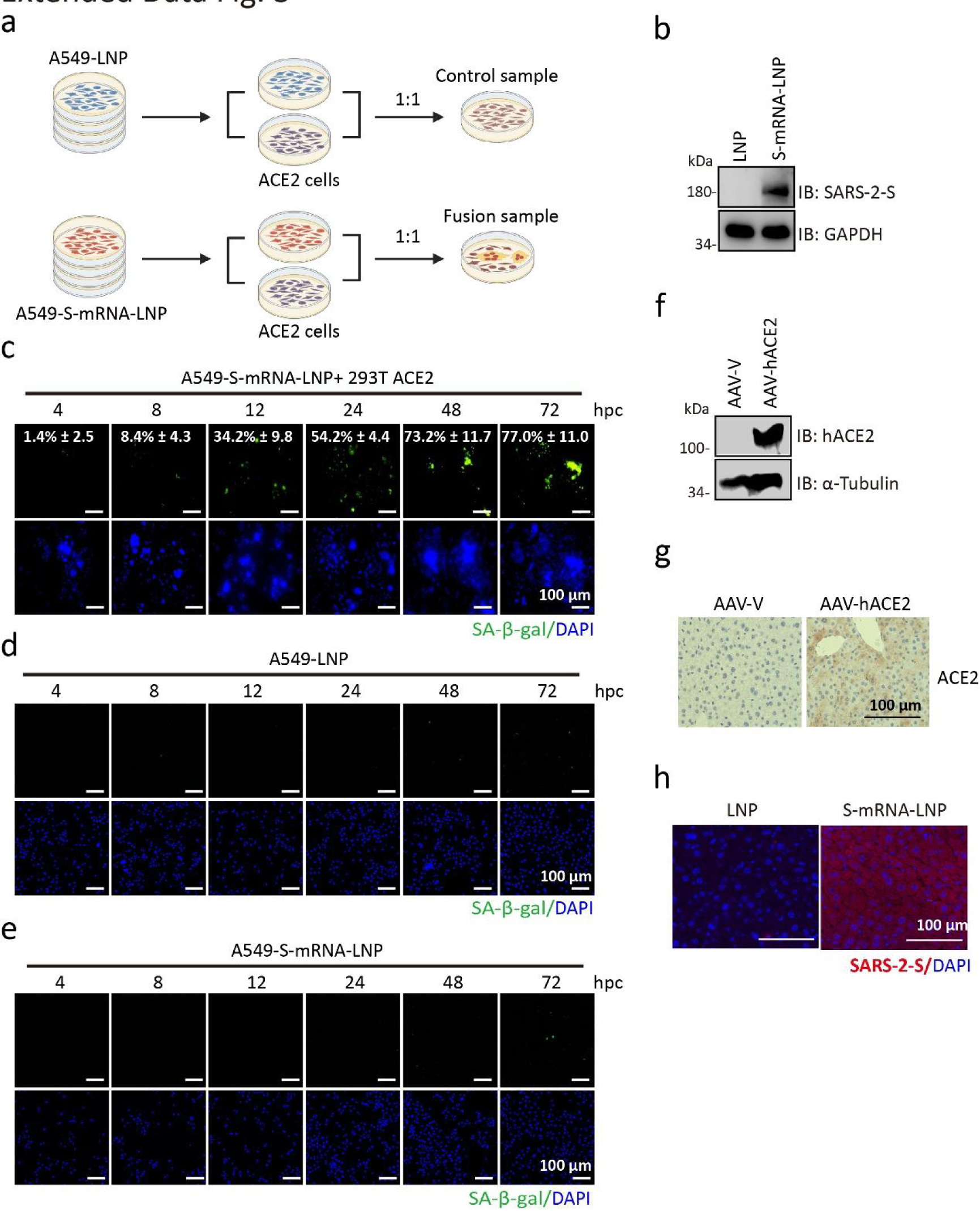
SARS-2-S delivered by mRNAs triggers syncytial senescence. a, Illustration of the cell coculture system mediated by S-mRNAs. Cells were treated with S-mRNA-LNP for 24 h, followed by coculture with ACE2-expressing cells for the indicated times. A mixture of A549-LNP and A549-ACE2 cells was used as a nonfusion control. b, Immunoblot analysis of S expression in A549 cells treated with S-mRNA-LNP. GAPDH was used as a loading control. c, SA-β-gal staining of A549-S-mRNA-LNP cells cocultured with 293T-ACE2 cells for the indicated times. Green, SA-β-gal staining; blue, nuclei DAPI staining. Scale bars represent 100 μm. d, e, SA-β-gal staining of LNP-(d) or S-mRNA-LNP-treated (e) A549 cells cultured for the indicated times. Green, SA-β-gal staining; blue, nuclei DAPI staining. Scale bars represent 100 μm. f, g, h ACE2 expression or S expression (h) in livers from mice injected with S-mRNA-LNPs or LNPs by immunoblotting (f) or immunohistochemical staining (g). All images are representative of three independent experiments.

**Fig. S4.**
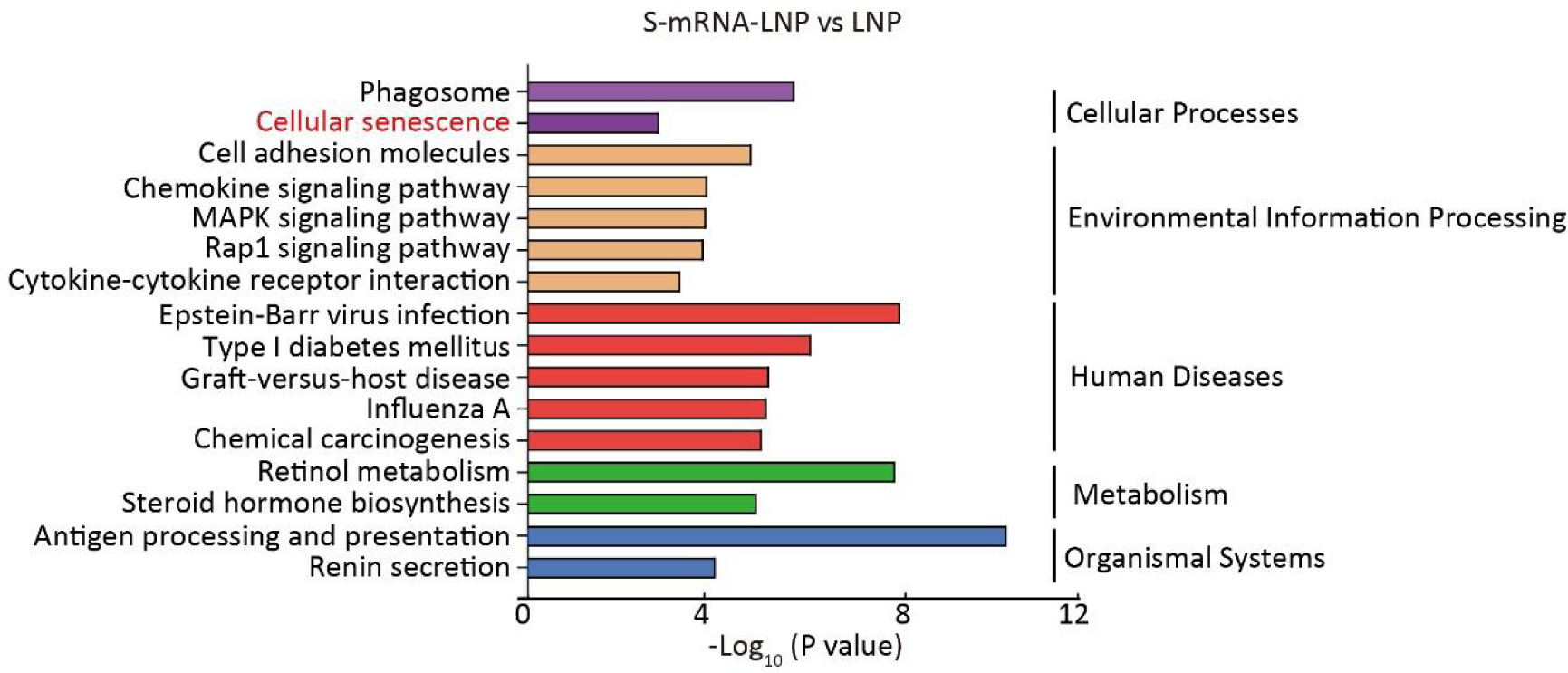
SARS-2-S delivered by mRNAs triggers syncytial senescence. Pathways enriched in DEGs identified in the livers of mice injected with LNPs or S-mRNA-LNPs according to KEGG functional pathways at level 1. The bar plot shows significantly dysregulated pathways (P < 0.05), with Fisher’s exact test P values shown on the x-axis.

**Fig. S5.**
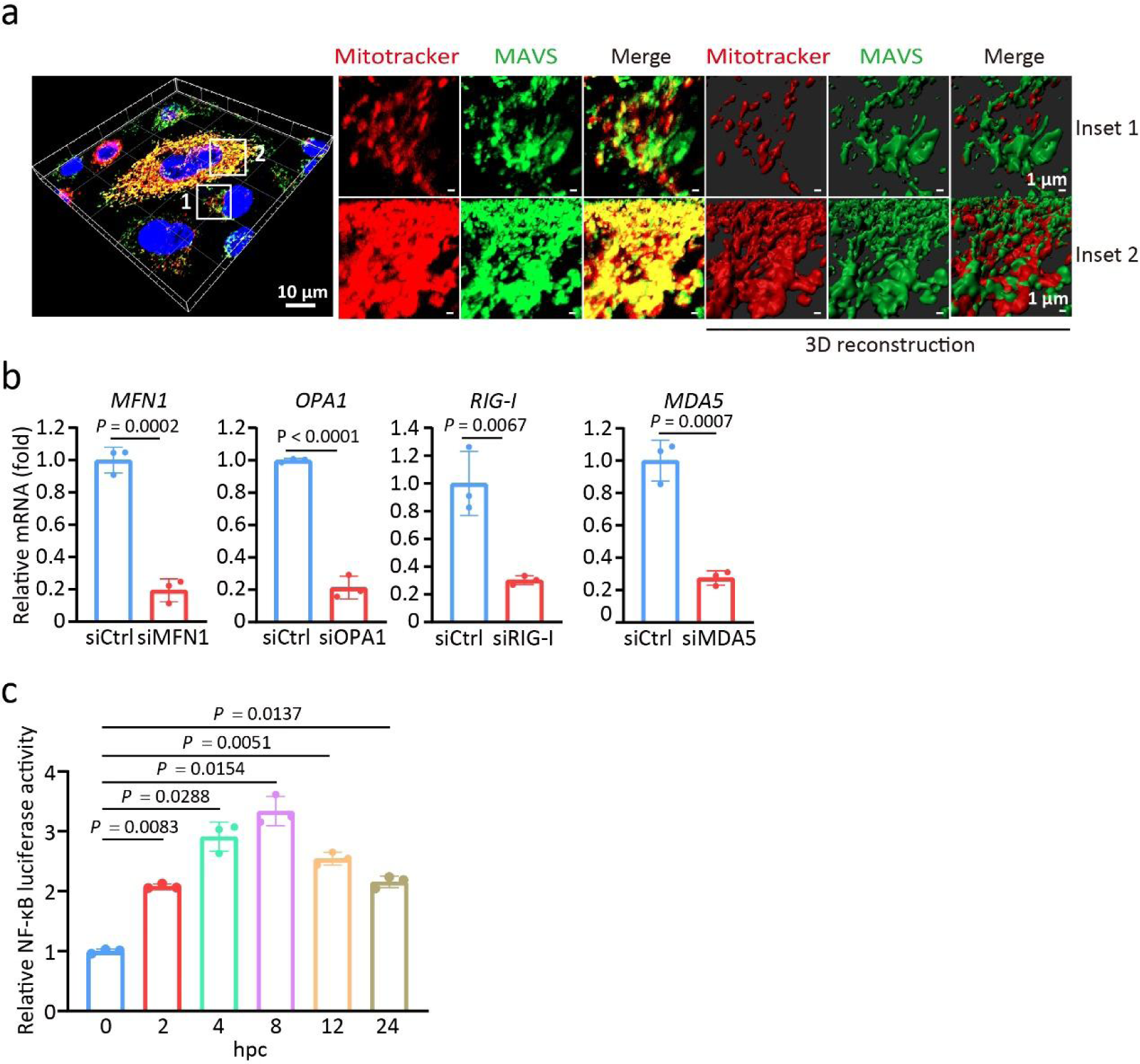
SARS-2-S syncytia induce the formation of functional MAVS aggregates dependent on RIG-I. a, Confocal microscopic images of MAVS and mitochondria in cocultured A549 cells at 4 hpc. Red, MitoTracker staining for mitochondria; green, MAVS staining; blue, DAPI staining. A magnified view of the boxed region is shown in the middle panel, and a 3D reconstruction of the Z stacks is shown in the right panel. Scale bars represent 10 μm (whole image) and 1 μm (magnified). b, Knockdown efficiency of siMAVS, siRIG-I, siMFN1, or siOPA1 compared to siCtrl by RT-qPCR. c, Relative luciferase activity of NF-κ B in cocultured A549 cells for the indicated times. Luciferase activity was normalized to the values at 0 hpc. All quantified data in this figure are shown as the means ± SDs of n = 3 independent experiments. Statistical significance was determined with two-tailed Student’s t test (b), one-way ANOVA and Bonferroni’s post hoc analysis (c).

**Fig. S6.**
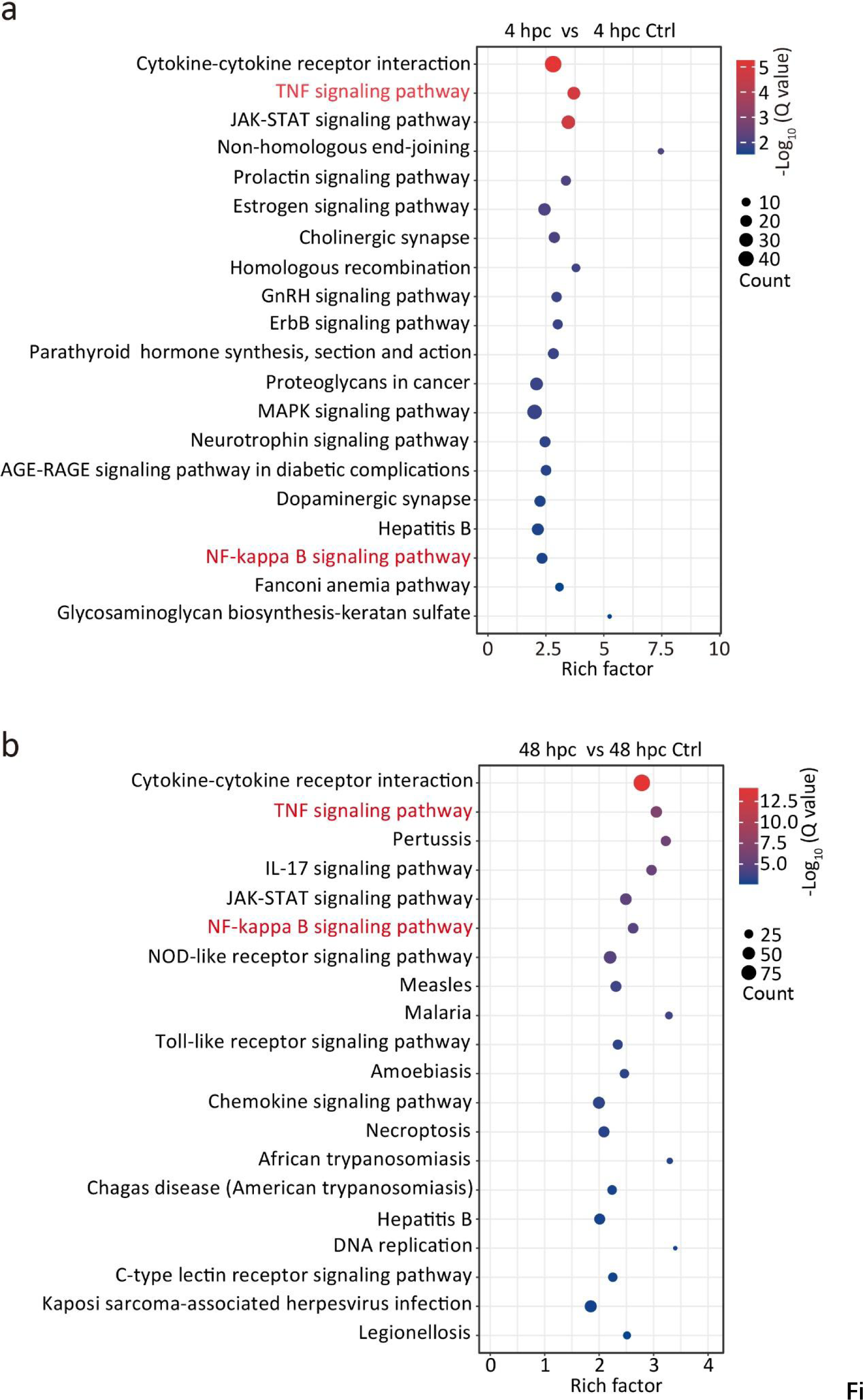
MAVS-TNFα regulates the survival and senescence of SARS-2-S syncytia. a, b, Twenty most significantly enriched pathways in cocultured A549 cells at 4 hpc (a) and 48 hpc (b) according to KEGG pathway analysis. The enriched terms are shown on the y-axis, and the P values (log transformed) assessing significant enrichment are shown on the x-axis with Fisher’s exact test. The enrichment degree of KEGG was measured by enrichment factors (rich factor), P value and the number of genes enriched in this pathway.

**Fig. S7.**
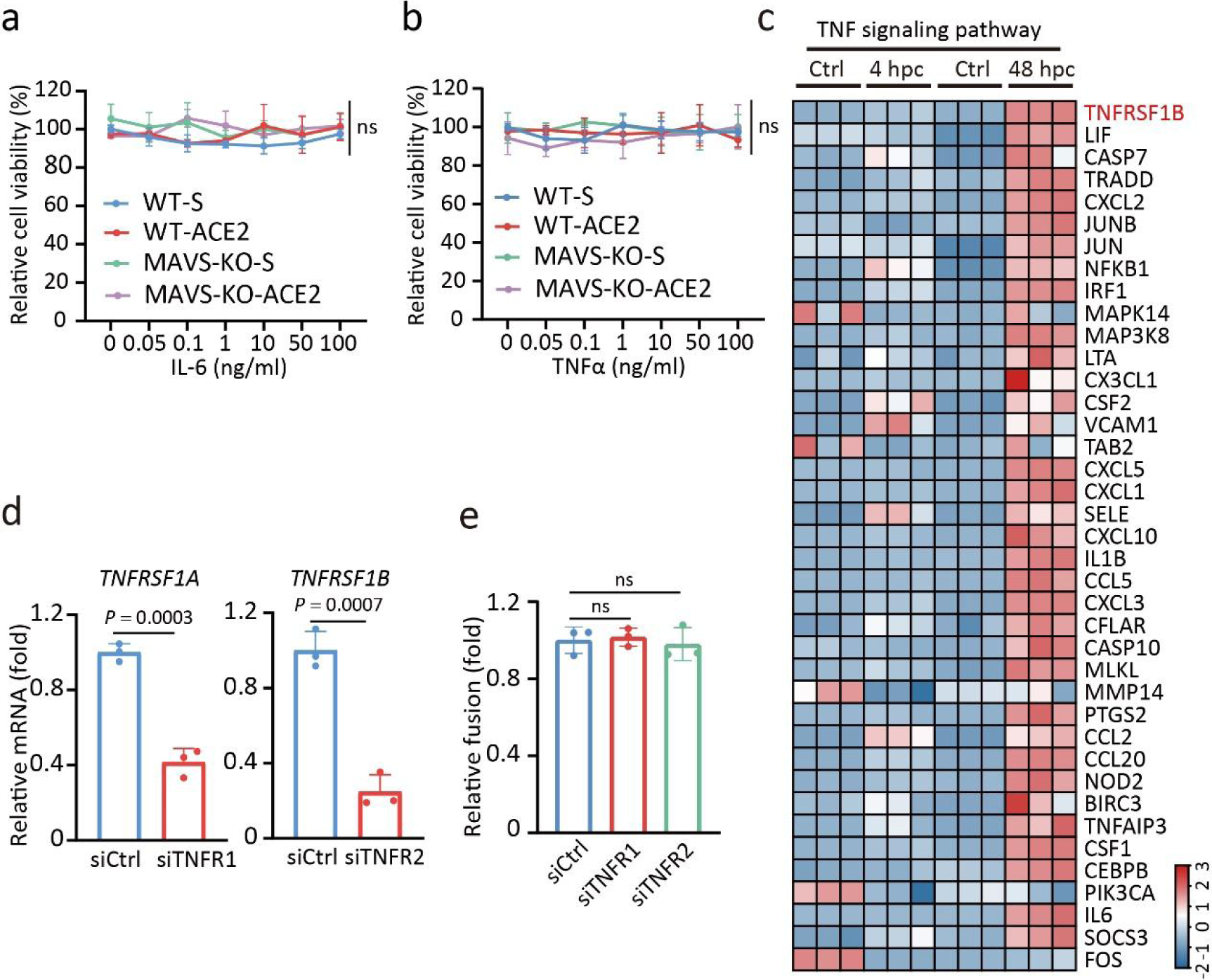
MAVS-TNFα regulates the survival of SARS-2-S syncytia. a, b, Relative viability of WT-S, WT-ACE2, MAVS-KO-S, and MAVS-KO-ACE2 cells treated with IL-6 (a) or TNFα (b) at the indicated concentrations. The cell viability of cells treated with PBS was set to 100%. c, Heatmap of the critical DEGs of the TNF signalling pathway in cocultured A549 cells at 4 hpc and 48 hpc. d, Knockdown efficiency of siTNFRSF1A or siTNFRSF1B compared to siCtrl by RT-qPCR. e, Relative fusion of siTNFR1 and siTNFR2 cocultured A549 cells at 24 hpc by β-galactosidase assay. The fusion level of siCtrl cocultured cells at 24 hpc was set to 1. All quantified data are shown as the means ± SDs of n = 3 independent experiments. Statistical significance was determined with two-tailed Student’s t test (d), one-way ANOVA and Bonferroni’s post hoc analysis (a, b, e).

**Fig. S8.**
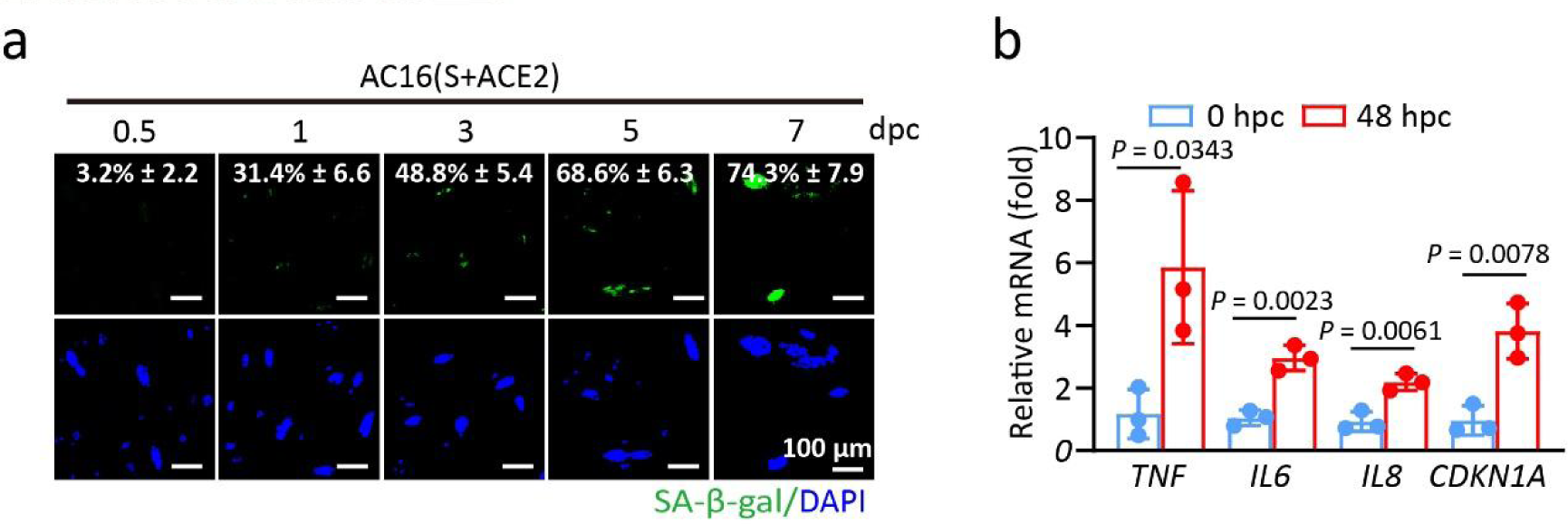
Senescent SARS-2-S syncytia shrinkage triggers WNK1 phase-separation. a, SA-β-gal staining of AC16 cells cocultured for the indicated days (dpc). Scale bars represent 100 μm. Green, SA-β-gal staining; blue, nuclear DAPI staining. b, Normalized expression of *TNF*,*IL6,IL8,andCDKN1A* in AC16 cells at 48 hpc from a relative to that at 0 hpc by RT-qPCR. All quantified data are shown as the mean ± SD of n = 3 independent experiments. Statistical significance was determined using two-tailed Student’s t test (b).

**Fig. S9.**
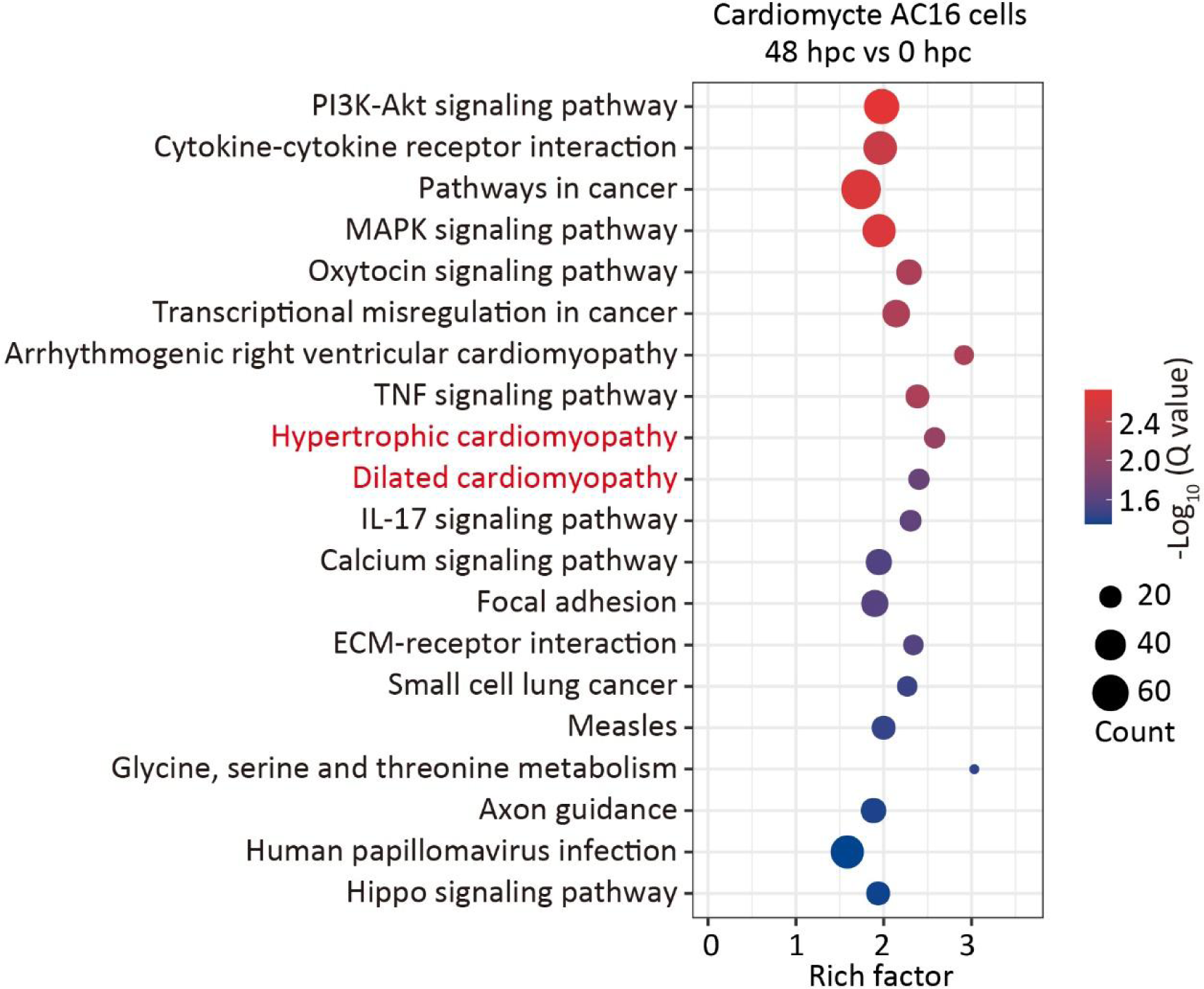
WNK1 activation in senescent SARS-2-S syncytia impairs cardiac metabolism. Twenty most significantly enriched pathways in cocultured AC16 cells at 48 hpc versus 0 hpc according to KEGG pathway analysis. The enriched terms are shown on the y-axis, and the P values (log transformed) assessing significant enrichment are shown on the x-axis with Fisher’s exact test. The enrichment degree of KEGG was measured by enrichment factors (rich factor), P value and the number of genes enriched in this pathway.

**Fig. S10.**
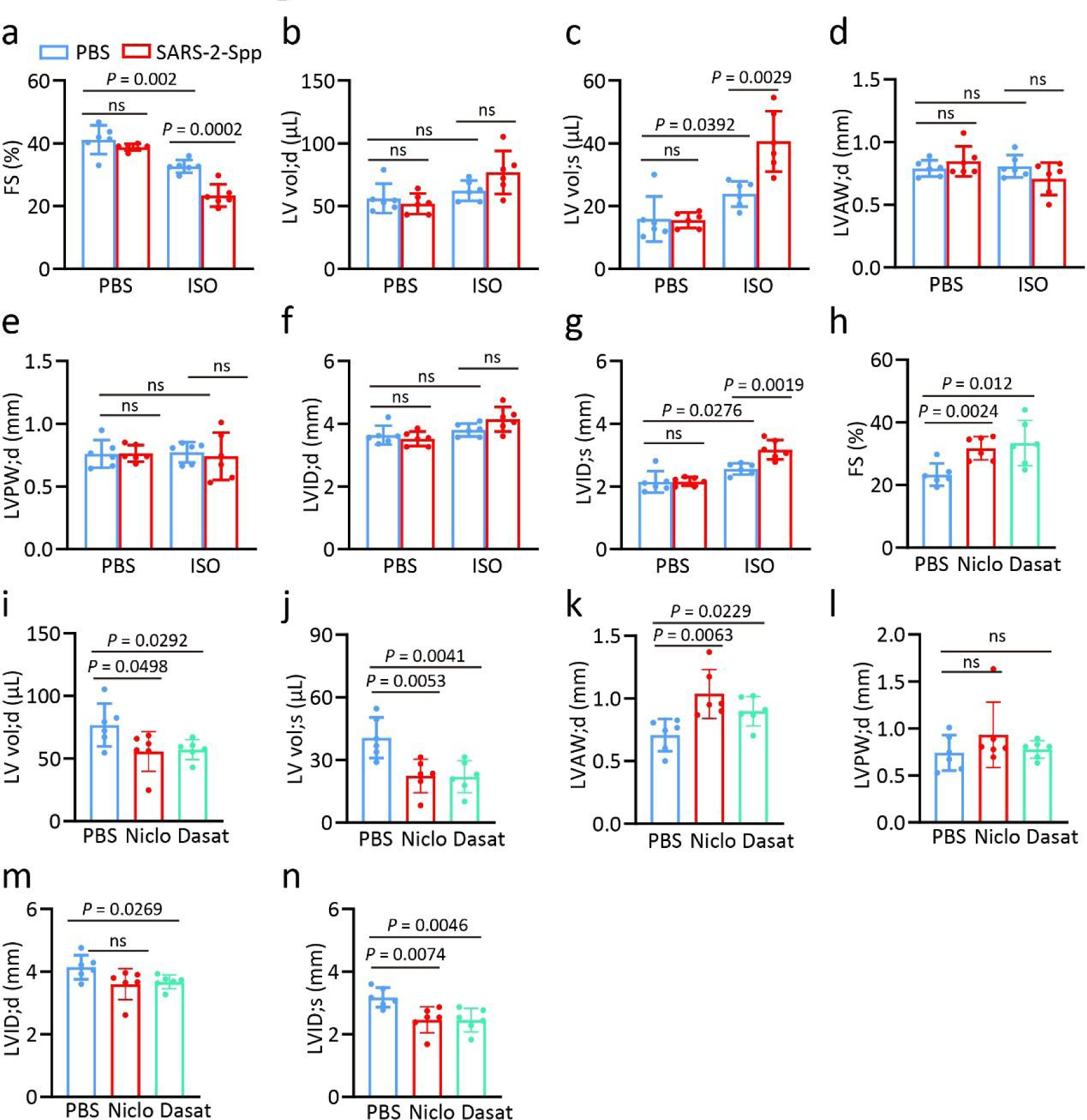
Senescent SARS-2-S syncytia exacerbated heart failure. a-g, Fractional shortening (FS) (a), LV end-systolic volumes (LV Vol;d) (b), end-diastolic volumes (LV Vol; s) (c), left ventricular end-diastolic anterior wall thickness (LVAW;d) (d), left ventricular posterior wall thickness (LVPW;d) (e), left ventricular end-diastolic diameter (LVID;d) (f), and left ventricular end-systolic diameter (LVID;s) (g) of SARS-2-Spp-infected mice with or without ISO-injection. h-n, FS (h), LV Vol;d (i), LV Vol; s (j), LVAW;d (k), LVPW;d (l), LVID;d (m), and LVID;s (n) of SARS-2-Spp-infected ISO-mice treated with PBS, niclosamide (nic), or dasatinib (das). All quantified data are presented as the mean ± SD of n = 6 independent experiments. Statistical significance was determined with one-way ANOVA and Bonferroni’s post hoc analysis (a, b, c, d, e, f, g, h, i, j, k, l, m, n).

**Fig. S11.**
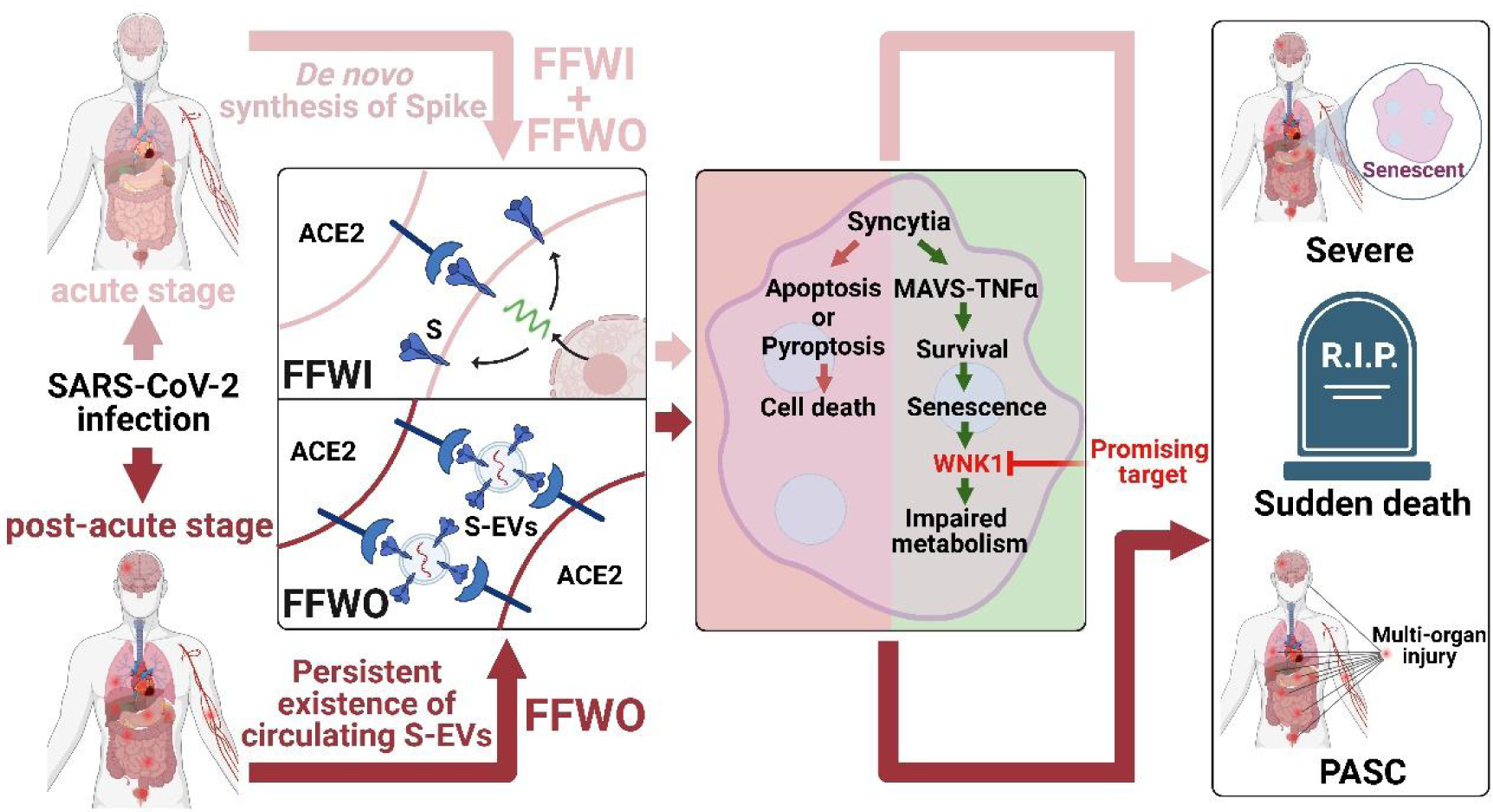
SARS-CoV-2 spike-induced syncytia are senescent and contribute to exacerbated heart failure. SARS-2-S induced syncytia via FFWI and FFWO manner exhibits a senescence-like phenotype regardless of the cell type or syncytium nucleus number. Mechanistically, RIG-I-MAVS drives the TNFα-dependent survival and senescence fate of SARS-2-S syncytia. The susceptibility of patients with heart failure to more pronounced complications during the acute and post-acute stages of SARS-CoV-2 infection could be related to the senescent outcome of SARS-2-S syncytia, and WNK1 inhibitor may be a druggable target in the management of heart failure associated with SARS-CoV-2 infection.

